# Reversing transgene silencing via targeted chromatin editing

**DOI:** 10.1101/2025.10.28.685244

**Authors:** Elia Salibi, Sebastian Palacios, Eric Lu, Ron Weiss, Thorsten M. Schlaeger, James J. Collins, Domitilla Del Vecchio

**Affiliations:** Department of Mechanical Engineering, Massachusetts Institute of Technology, Cambridge, MA, 02139, USA; Institute for Medical Engineering and Science, Massachusetts Institute of Technology, Cambridge, MA, 02139, USA; Department of Biological Engineering, Massachusetts Institute of Technology, Cambridge, MA, 02139, USA; Boston Children’s Hospital Stem Cell Program, Boston Children’s Hospital, Boston, MA, 02115, USA; Broad Institute of MIT and Harvard, Cambridge, MA, USA; Wyss Institute for Biologically Inspired Engineering, Harvard University, Boston, MA, USA

**Keywords:** CHO-K1, human stem cells, genome engineering, epigenetics, synthetic biology

## Abstract

Advances in cell engineering and mammalian synthetic biology are enabling the development of genetically engineered cells to uncover biological principles and develop next-generation biotechnologies. However, epigenetic silencing of transgenes remains a major challenge in the field, particularly for human induced pluripotent stem cells (hiPSCs). Here, we employ targeted chromatin editing to study the dynamics of transgene silencing at integrated reporters in both CHO-K1 cells and hiPSCs. We first induce silencing by transiently recruiting writers of DNA methylation or histone 3 lysine 9 trimethylation (H3K9me3) to educate a chromatin modification model, which predicts DNA demethylation is sufficient for transgene reactivation. Accordingly, we demonstrate efficient and stable reactivation in both cell types through transient recruitment of a chromatin regulator that erases DNA methylation, irrespective of whether durable silencing was achieved with DNA methylation, H3K9me3 or triggered by endogenous cellular machinery. These results shed light on molecular mechanisms at play in transgene silencing and provide engineering tools for potent and specific reactivation in mammalian cells.

## Introduction

The ability to engineer cell function is a cornerstone of biological research, engineering, and medicine ^1–4^. In particular, mammalian synthetic biology allows the study of cell physiology in normal and diseased states with unprecedented control ^5^, enables the manufacture of effective biotherapeutics ^6,7^, and supports the development of next-generation cellular therapies ^8–11^. However, a major obstacle hampering the progression of such technologies from bench to bedside is epigenetic silencing of transgenes, the loss of gene expression in genomic integrations over time that is not caused by changes in the DNA sequence ^12–21^.

Although transgene silencing has been observed across various cell types and species, it has high relevance in CHO cells, commonly used for industrial protein production, as well as in human induced pluripotent stem cells (hiPSCs) for clinical applications. The production of therapeutic proteins in CHO cells has facilitated the diagnosis and treatment of diseases ranging from autoimmunity to cancer ^22^. Despite optimizations in construct design, culture conditions and cell lines, there remains obstacles, particularly for difficult to produce antibodies ^23–27^. In this case, epigenetic silencing diminishes the long-term expression profiles of genetically engineered CHO cells ^28–32^. Similarly, hiPSCs derived from somatic cells have immense potential due to their ability to differentiate into any cell type. As such, these cells hold unique promise to advance the fields of human development and medicine. While human stem cells can be guided to desired cell states with chemical compounds ^33–36^, advances in genome engineering and synthetic biology are enabling genetic programming of hiPSCs by intrinsically controlling their differentiation to specific cell types in space and time ^37–40^. However, epigenetic silencing of transgenes poses a significant efficacy and safety risk, undermining stem cell-derived therapies and delaying clinical applications. Therefore, strategies to prevent or reverse epigenetic silencing are needed.

Epigenome engineering has emerged as a powerful tool for precise gene regulation by targeted chromatin remodeling ^41–44^. The regulators comprise a DNA binding domain for specific sequence targeting and an effector domain to either directly add/remove chemical modifications to chromatin or serve as a docking site for the recruitment of endogenous factors ^45–48^. The field has largely focused on the activation and silencing of endogenous genes to treat diseases associated with aberrant gene expression ^49–57^. For instance, dCas9 fused to DNMT3A/L and KRAB domains was used to downregulate *pcsk9* gene expression with the aim of decreasing circulating low-density lipoprotein cholesterol levels *in vivo* ^58^. By fusing DNMT3L cofactor to a DNA binding domain and a histone 3 tail, it was possible to recruit and activate endogenous DNMT3A in cells to methylate and repress a locus involved in prion protein production ^59^. Another study recruited dCas9-TET1 fusion protein to specifically demethylate the *brca1* tumor suppressor gene implicated in breast cancer ^60^. Moreover, epigenome editing was used to rescue diseased phenotypes in neurons derived from Rett Syndrome and Fragile X Syndrome patients by TET1-mediated DNA demethylation at *mecp2* and *fmr1* loci, respectively ^61,62^. DNA demethylating systems have further been used to reverse the silencing of the imprinted locus involved in Prader-Willi syndrome ^63^. By combining orthologous dCas9 proteins for simultaneous DNA methylation and demethylation at different loci, it was further possible to alter glycosylation patterns of BG1 cells by targeting *hnf1a* and *mgat3* genes ^64^.

In contrast to epigenetic editing of endogenous genes, epigenetic editing of transgene loci remains underexplored, especially in hiPSCs. In CHO-K1 cells, silencing dynamics of a reporter gene following epigenetic editing with different chromatin regulators was studied in a human artificial chromosome ^46^ and in the natural chromosome ^65^. In K562 cells, endogenous DNMT3A recruited to a methylation sensitive reporter by rTetR-DNMT3L led to durable silencing ^66^. The synergy between writers of DNA methylation and H3K9me3 was studied by recruiting combinations of epigenetic regulators (KRAB/DNMT3A/L) in multiple cell types ^45,47,49,67–70^. The role of the CpG content on the silencing rate of a reporter gene was further investigated in CHO-K1 cells by building a synthetic promoter library with varying CpG content ^71^. Also using a reporter transgene, a different study characterized DNMT3A/L variants for their methyltransferase and gene silencing activities in HEK293 and Jurkat T cells ^72^. Few studies have characterized reactivation dynamics of a reporter transgene ^45,70,73,74^. These studies either fused the TET1 catalytic domain or different epigenetic activators to dCas9 in HEK393T cells ^45,70^ or TET1 to the rTetR-transactivator (rtTA) in mouse embryonic stem cells ^73,74^. While the above studies provide significant advancements, they are performed in different cellular and genomic contexts using different integration methods and promoters, confounding the rules and molecular mechanisms that govern transgene silencing and reactivation.

Here, we explore the use engineered chromatin regulators to silence reporters chromosomally-integrated at a specific safe harbor locus in both CHO-K1 cells and hiPSCs to define the roles of H3K9me3 and DNA methylation in silencing with the aim of identifying methods to reverse endogenous epigenetic silencing. We first characterized gene expression and chromatin states in CHO-K1 cells following induction of H3K9me3 or DNA methylation by transient transfection of KRAB or DNMT3A domains, respectively. We then repeat the experiments using an identical integrated reporter system in hiPSCs and observed different silencing dynamics after transient KRAB recruitment. We used the data to educate a chemical reaction model of chromatin modifications, which suggests a secondary role for H3K9me3 during silencing. Based on these predictions, we reactivate DNMT3A- and KRAB-silenced transgenes in hiPSCs by TET1-mediated DNA demethylation alone, and further demonstrate successful reporter reactivation following endogenous epigenetic silencing. Finally, we characterize silencing and reactivation dynamics of reporters randomly integrated into the genome by viral transduction, obtaining results that largely mirror those observed with site-specific chromosomal integration of the reporter.

## Results

### Control of epigenetic states by targeted chromatin editing in somatic mammalian cells

We first set out to investigate whether we could control epigenetic states in chromosomal integrations via chromatin editing. To this end, we utilized an experimental system that we previously developed in CHO-K1 cells (Fig. 1A), a model cell line for synthetic biology with wide use in biotechnology ^65^. This system comprises a single-copy, site-specific integration of a reporter cassette in the Rosa26 locus. The reporter cassette includes a fluorescent protein (EBFP2) driven by a constitutive promoter (hEF1a) with upstream operator sites that allow recruitment of DNA binding domains fused to chromatin regulators (Fig. S1A), thus enabling targeted editing of chromatin at the locus (See Methods). To assess our ability to control chromatin states, we subjected cells in the active state to transient transfection with DNMT3A, a *de novo* DNA methylation writer ^75^, fused to rTetR for sequence-specific targeting (rTetR-DNMT3A). We then transiently transfected silenced cells with rTeTR-TET1, comprising the TET1 catalytic domain ^57^ fused to rTetR (or TetR when specified) for sequence-specific targeting (see Methods and Supplementary Table 1). Reporter gene expression was monitored by time-resolved flow cytometry (Fig. 1B).

**Fig. 1.**
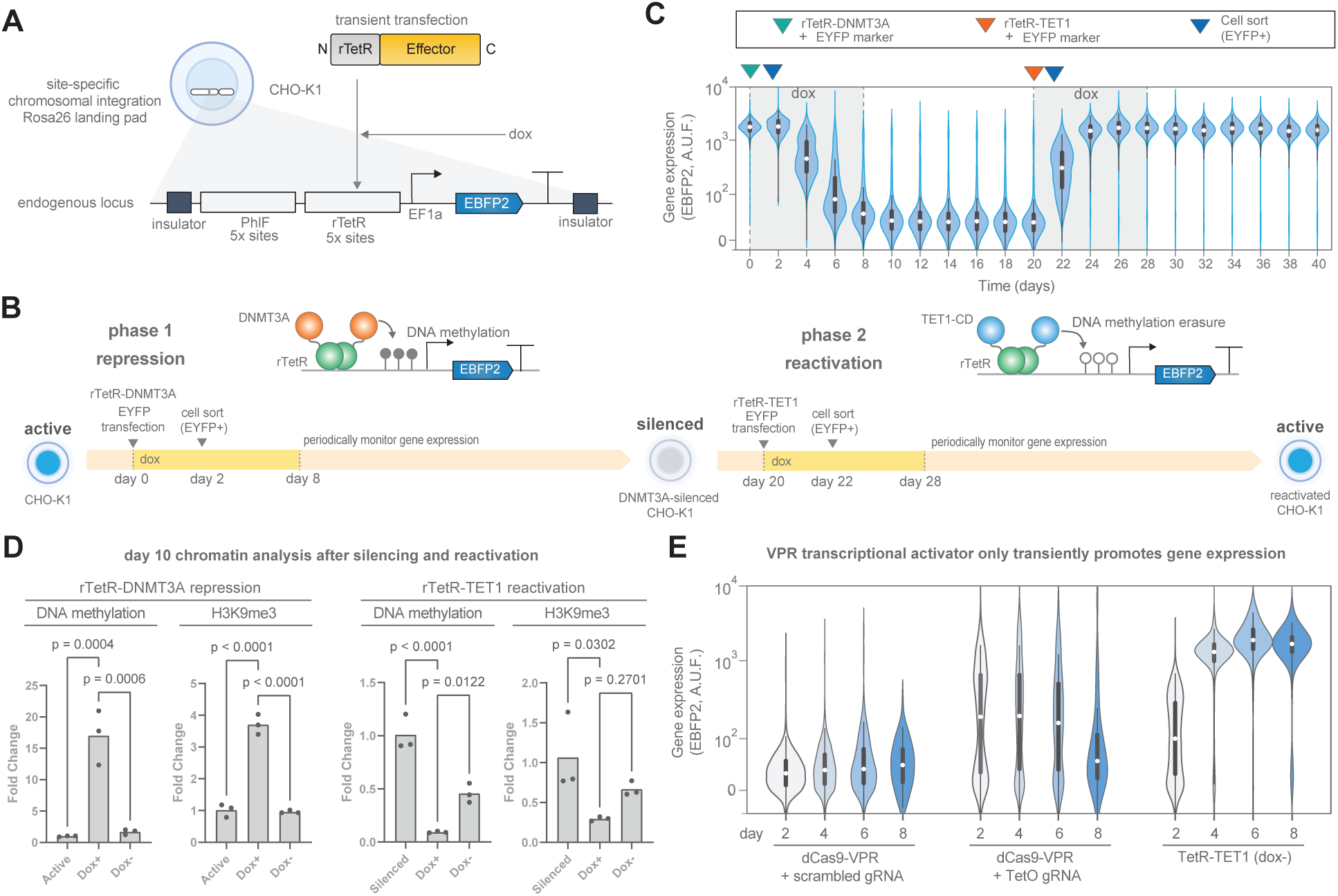
Targeted chromatin editing enables reversible control of epigenetic and transcriptional state in adult mammalian cells. **A.** Schematic of the experimental system. The reporter gene is integrated in the ROSA26 locus in CHO-K1 cells. The promoter (hEF1a core + intron) drives expression of EBFP2 and is downstream of binding sites for PhlF and rTetR. The construct is flanked by cHS4 chromatin insulators. ’dox’ is doxycycline, a small molecule that enables rTetR operator binding. **B.** Schematic of the experimental workflow for targeted chromatin editing in the engineered CHO-K1 cells. **C.** Violin plots of flow cytometry measurements during CHO-K1 cell state switching by sequential transient rTetR-DNMT3A and rTetR-TET1 transfections. Data shown are from one replicate from three independent replicates. **D.** Quantitative PCR analysis after methylated DNA immunoprecipitation (MeDIP) or CUT&RUN targeting H3K9me3. Bars represent mean values of three independent replicates. Statistics were performed with an ordinary one-way ANOVA with Dunnet correction for multiple comparisons. **E.** Violin plots of flow cytometry measurements after recruitment of dCas9-VPR or TetR-TET1 over the course of 8 days. Data are plotted from three independent replicates. In all violin plots, the white dot represents the median, the thick box represents the interquartile range (IQR) and the thin gray line represents

We co-transfected rTetR-DNMT3A along with EYFP, which we use as a proxy for rTetR-DNMT3A expression levels ^76,77^, isolated transfected CHO-K1 cells (EYFP+) by fluorescence-activated cell sorting (FACS) two days later and monitored fluorescence changes by flow cytometry. We observed a steady decrease in reporter gene expression over the course of 8 days, reaching complete silencing by day 10 (Fig. 1C). Gene expression levels remained low after withdrawal of doxycycline (dox), indicating that the reporter was stably silenced. After 20 days, we co-transfected the silenced cells with rTetR-TET1 and EYFP transfection marker, isolated transfected cells (EYFP+) by FACS and continued monitoring fluorescence. Already 4 days post rTetR-TET1 transfection, the reporter was fully reactivated and remained active after dox withdrawal, indicating stable reactivation. We additionally compared reactivation strength of different transfection levels of rTetR-TET1 by FACS, noting that medium to high levels displayed the best performance in terms of trade-off between reactivation strength and leakiness (Fig. S1B-C, Bin 2). At very high levels (S1B-C, Bin 3), we observed significant reactivation even in the absence of dox, suggesting leaky rTetR binding. We then asked whether it was possible to reactivate gene expression after the cells had been cultured in the silenced state for over 4 weeks (S1D). Indeed, the cells displayed strong and efficient reactivation when TET1 was targeted, while mock plasmid or a catalytically dead TET1 domain showed no response, indicating that time spent in the silenced state does not affect reactivation efficiency and that TET1 catalytic activity is necessary and sufficient for reactivation. While DNMT3A recruitment durably silenced transgene expression, transient transfections with KRAB domain, a regulator known to indirectly recruit histone 3 lysine 9 trimethylation (H3K9me3) ^78^, led to transient repression in CHO-K1 cells that lasted only while KRAB was detectable (Fig. S1E). We and others have previously shown that no DNA methyaltion is deposited in CHO-K1 following targeted KRAB treatment, hinting at a potential mechanism for durable silencing ^46,65^. We observed no silencing when transfecting the KRAB or DNMT3A domains without targeting (Fig. S2A), and little to no reactivation when transfecting TET1 or rTetR alone (Fig. S2B), thus ensuring that both DNA binding and catalytic activity are required. These results demonstrate the ability of our system to be toggled between active and silenced states efficiently following transient transfection of the engineered chromatin regulators.

We further characterized the chromatin marks at the reporter locus 10 days after transfection with the chromatin regulators. We performed quantitative PCR (qPCR) following methylated DNA immunoprecipitation (MeDIP-qPCR) to assess DNA methylation levels and Cleavage Under Targets and Release Using Nuclease (CUT&RUN-qPCR) to assess H3K9me3 levels (see Methods). The data show that DNMT3A recruitment results in enrichment for DNA methylation as expected with a 15-fold increase compared to active cells (Fig. 1D, left). We also detected an increase of H3K9me3 although to a lesser degree, suggesting that DNA methylation recruits writers of H3K9me3, which is consistent with prior reports ^46^. In contrast, we and others have shown that KRAB recruitment in CHO-K1 cells results in H3K9me3 enrichment without DNA methylation ^46,65^. After targeted chromatin editing with rTetR-TET1, both DNA methylation and H3K9me3 marks were reduced to approximately 10% and 25% of that in silenced cells, respectively (Fig. 1D, right), suggesting that H3K9me3 cannot sustain itself in the absence of DNA methylation.

In addition to employing the TET1 catalytic domain, we attempted to reactivate the silenced transgene with other relevant chromatin regulators (Fig. S2C). In particular, we employed catalytic domains of DOT1L, a histone 3 lysine 79 methyltransferase (H3K79me2), the p300 core domain, a histone 3 lysine 27 acetyltransferase (H3K27ac), and PRDM9, a histone 3 lysine 4 methyltransferase (H3K4me3), and their combinations since H3K79 and H3K4 methylation were described to act synergistically ^48,54,79^. No reporter gene expression could be detected by flow cytometry for those conditions that excluded the TET1 domain. Nevertheless, it does not rule out the possibility that the protein fusions might be misfolding or that recruiting these marks upstream the promoter is not the optimal location. We can conclude, however, that recruiting TET1 is sufficient to rescue expression from a transgene loaded with DNA methylation. To test whether transcriptional activator domains would promote gene expression, we utilized the VP64-p65-Rta (VPR) transcriptional activator ^80^ fused to dCas9 with a guide RNA (gRNA) targeting all the TetO sites (Fig. 1E, S2D). VPR recruitment resulted in a burst of gene expression soon after transfection but slowly returned to basal levels over the course of 8 days. In contrast, TET1 recruitment resulted in a steady increase in gene expression that remained stable for the duration of the experiment, again highlighting the need to demethylate the locus for durable reactivation. We could recapitulate the results when swapping the hEF1a promoter with other frequently used constitutive promoters (CAG, PGK and UbC, see Fig. S3A-B and Supplementary Table 2), showing responsiveness to both silencing by rTetR-DNMT3A and reactivation by rTetR-TET1 (Fig. S3C-D). Taken together, these data suggest that DNA methylation has a primary role in transgene silencing due to its long-term stability while H3K9me3 alone cannot sustain itself and hence plays a secondary role in epigenetic silencing where DNA methylation is possible.

### Dissecting silencing dynamics in hiPSCs using targeted chromatin editing

We extended our investigation to hiPSCs to characterize the interactions between DNA methylation and H3K9me3 during silencing. To this end, we sought to construct the reporter gene in human stem cells utilizing landing pad technology for site specific recombinase mediated integration ^81,82^, which could closely recapitulate the chromosomal integration of the reporter gene we developed in CHO-K1 cells. We consequently developed an hiPSC landing pad cell line and performed chromosomal integration of the reporter system in human stem cells (Fig. 2A, S4A and Methods). The chromosomal integration lies in the AAVS1 locus, a genomic region that is widely utilized for transgene integration in human cells ^83^. Despite its widespread adoption, there are mixed reports of transgene expression stability in the AAVS1 locus in human stem cells ^20,84–87^, hinting at a context-dependent behavior, potentially stemming from the insulator-promoter-gene triad employed.

**Fig. 2.**
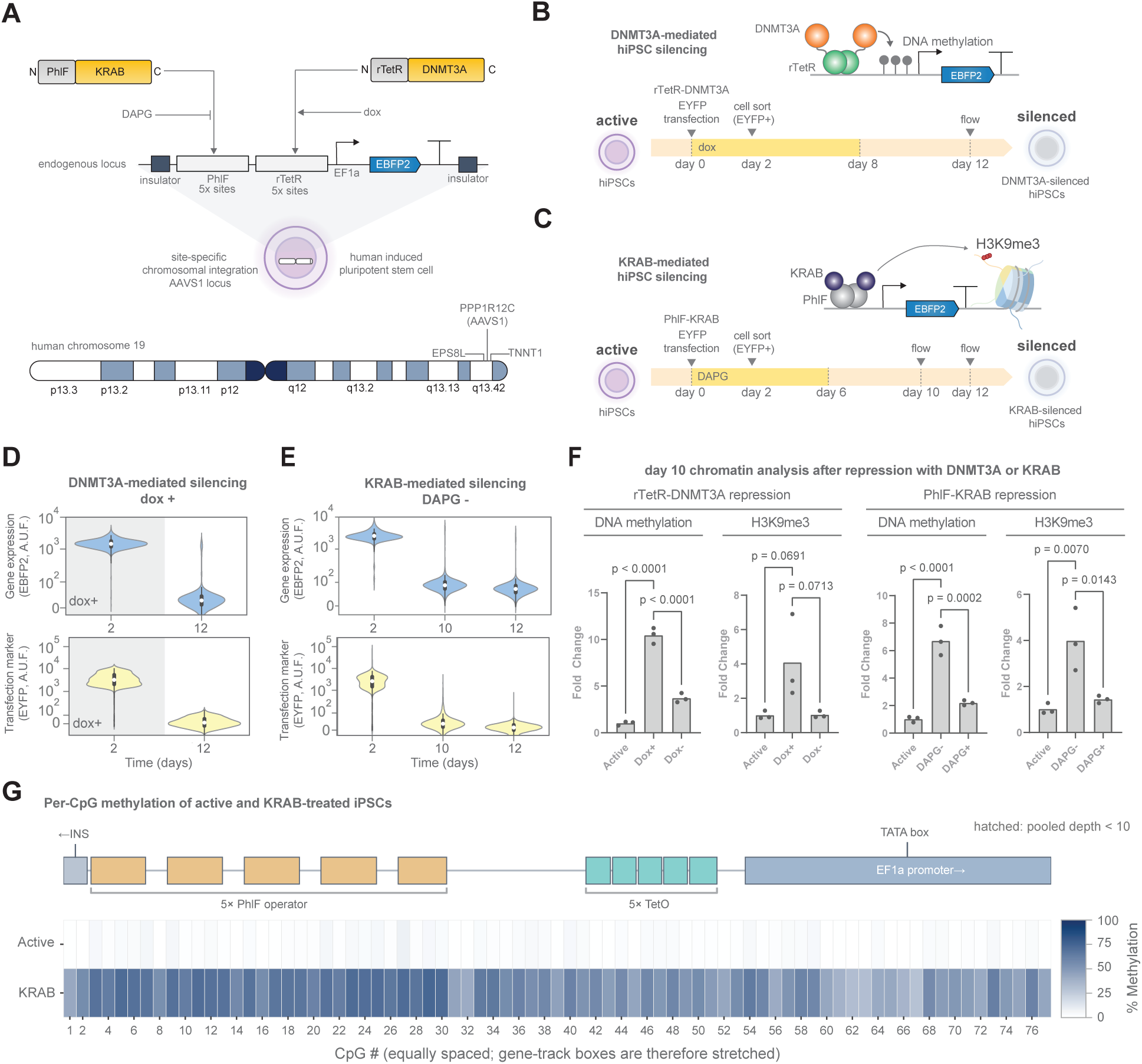
**Different engineered chromatin regulators enable stable silencing in human induced pluripotent stem cells**. **A.** Schematic of the experimental system. The reporter system is identical to that used in CHO-K1 cells except that the integration site is the AAVS1 locus in hiPSCs (see Methods and Fig. S3A). ’DAPG’ is 2,4-diacetylphloroglucinol, a small molecule that inhibits PhlF binding to its operator. **B-C.** Schematic of the experimental workflow for chromatin editing with **(B)** rTetR-DNMT3A or with **(C)** PhlF-KRAB. **D-E.** Violin plots of flow cytometry measurements after transient **(D)** rTetR-DNMT3A or **(E)** PhlF-KRAB recruitment. Data are plotted from three independent replicates. The white dot represents the median, the thick box represents the interquartile range (IQR) and the thin gray line represents 1.5x the IQR. **F.** Quantitative PCR analysis of chromatin after methylated DNA immunoprecipitation (MeDIP) or CUT&RUN targeting H3K9me3. Bars represent mean values of three independent replicates. Statistics were performed with an ordinary one-way ANOVA with Dunnet correction for multiple comparisons. **G.** Heatmap of targeted TAPS+ sequencing of the reporter transgene promoter. Methylation level of each CpG residue within the operator and promoter regions are shown for active and KRAB-silenced cells. Data reported are average from three independent replicates.

We set out to study different modalities of epigenetic silencing in human stem cells. Specifically, we aimed to silence gene expression via *de novo* DNA methylation or via recruitment of H3K9me3 writers using our engineered chromatin regulators (see Methods), rTetR-DNMT3A and PhlF-KRAB, respectively (Fig. 2B-C). To investigate silencing via *de novo* DNA methylation, we co-transfected rTetR-DNMT3A and EYFP transfection marker, isolated transfected stem cells (EYFP+) by FACS and performed flow cytometry measurements on day 2 just after sorting and on day 12 (Fig. 2B). The cells cultured with dox for 8 days exhibited efficient silencing at day 12, long after dox was omitted and the EYFP transfection marker was diluted indicating absence of rTetR-DNMT3A in the system (Fig. 2D). Thus, by transiently recruiting solely DNMT3A and depositing DNA methylation *de novo*, we achieved silencing that persisted even after withdrawal of the regulator. Next, we attempted to silence the cells via H3K9me3 deposition. We transfected the cells with PhlF-KRAB and EYFP, isolated transfected (EYFP+) stem cells using FACS and subsequently performed flow cytometry measurements at day 2, after sorting, and again at days 10 and 12 (Fig. 2C). The cells cultured in the absence of DAPG silenced efficiently and remained silenced after transient KRAB-mediated chromatin editing (Fig. 2E), indicating stable silencing similar to that achieved with DNMT3A, and contrasting with the transient silencing previously observed in CHO-K1 cells following KRAB treatment (Fig. S1E).

To shed light on how H3K9me3 recruitment led to stable silencing, we performed chromatin analysis of DNA methylation and H3K9me3 after recruitment of either mark. The data show that addition of one chromatin mark led to recruitment of the other in both cases (Fig. 2F). Specifically, DNMT3A recruitment resulted in enrichment for DNA methylation as expected with a 10-fold increase compared to active cells, and to a 4-fold increase of H3K9me3 compared to active cells (Fig. 2F, left). KRAB recruitment led to a 4-fold increase in H3K9me3 as expected, but also to a 6-fold increase in DNA methylation (Fig. 2F, left). We performed targeted methylated DNA sequencing at the transgene promoter region to corroborate our observations. Indeed, we found that KRAB-treatment led to an increase in the percentage of CpGs methylated within the sample population, ranging from 30-70% for each CpG residue, at all positions measured along the operator and promoter region up to the transcription start site (Fig. 2G and S4B).

To ensure that it is the H3K9me3 mark that recruits DNA methylation rather than the KRAB domain itself that is indirectly recruiting a *de novo* DNA methyltransferase, we tested different catalytic domains for histone methyltransferase activity by fusing them to TetR DNA binding domain. Specifically, we used catalytic domains of H3K9me3 writers SUV39 and G9A, as well as the H3K27 methyltransferase EZH2 as a control for histone methylation ^88^. We could only observe a stably silenced population with G9A catalytic domain after 10 days of treatment, achieving around 40% silenced cells (Fig. S5A). We further analyzed the chromatin marks of the silenced population after sorting and found a 10-fold increase in DNA methylation, but curiously, we could not detect a statistically significant fold-change increase in H3K9me3 mark (Fig. S5B). Targeted methyl CpG sequencing confirmed that 5-20% of each residue at the operator-promoter region in the population was, in fact, methylated (Fig. S5C-D).

Together, these data indicate that a positive feedback loop between H3K9me3 and DNA methylation exists in hiPSCs, wherein one mark recruits writers of the other. As a consequence, when transiently recruiting either DNMT3A or KRAB (or G9A) domain to the transgene locus in hiPSCs, we observed stable silencing and an increase in both DNA methylation and H3K9me3 marks, independent of the fusion protein employed for repression. Therefore, while DNA methylation mediates the recruitment of writers of H3K9me3 as in CHO-K1 cells (Fig 1D) ^46,65^, H3K9me3 mediates the recruitment of DNA methylation writers in hiPSCs but not in CHO-K1 cells ^65^, where H3K9me3 deposition results only in transient repression (Fig. S1E).

Intriguingly, when we repeated the silencing experiments by replacing the hEF1a promoter with the CAG promoter (we excluded UbC and PGK promoters due to low fluorescence signals from reporter hiPSC lines), DNMT3A alone had no effect on gene expression while KRAB resulted only in a transient repression (Fig. S6). However, co-transfection of both chromatin regulators led to stable silencing, indicating again a synergy between DNA methylation and H3K9me3, and suggesting that this synergy may depend on the specific promoter. The difference in behavior may stem from the variance in CpG number / density between the two promoters (see Fig. S3A), or from the CMV enhancer present in the CAG promoter.

### A chromatin modification model predicts silencing and reactivation dynamics

To dissect the molecular mechanisms at play during silencing across the two cell lines and promoters, we developed a chemical reaction model of DNA methylation and H3K9me3 based on a previously published model ^89^, but educated by our chromatin analysis data (Fig. 1D and Fig. 2F). After targeted chromatin editing with DNMT3A, we observed not only DNA methylation as expected, but also H3K9me3 (Fig. 1D and Fig. 2F, left), consistent with prior reports according to which methylated CpGs bind to proteins (Methyl-CpG-binding Proteins) that recruit H3K9me3 writers ^90^. The model thus includes the recruitment of H3K9me3 by DNA methylation. This leads to a chemical reaction path that goes from unmodified nucleosomes (U), through DNMT3A recruitment, to nucleosomes with DNA methylation only, denoted by y, to nucleosomes with both marks, denoted by z (Fig. 3A). Similarly, targeted chromatin editing with KRAB led to H3K9me3 deposition as expected, but also to DNA methylation in hiPSCs (Fig. 2F, right and 2G), consistent with prior reports indicating that H3K9me3 can be bound by proteins that recruit DNMT3A ^91,92^. Accordingly, the model accounts for the recruitment of DNA methylation by H3K9me3. This leads to a chemical reaction path from unmodified nucleosomes (U) to nucleosomes with H3K9me3 only (through KRAB recruitment), denoted by x, to nucleosomes z with both marks (Fig. 3A).

**Fig. 3.**
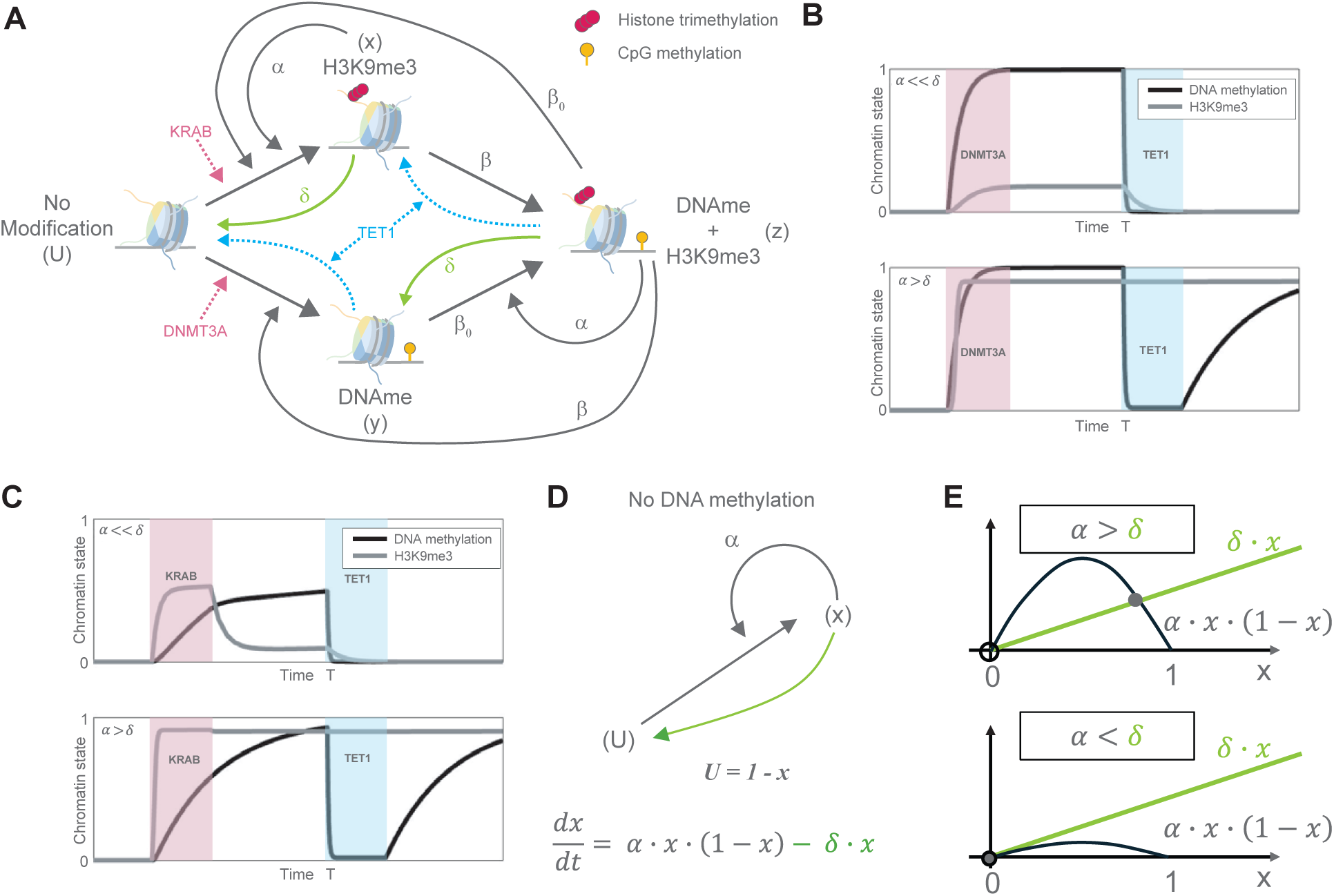
Model of the positive feedback between DNA methylation and H3K9me3 in hiPSCs. **A.** Straight arrows depict methylation reactions catalyzed by exogenous chromatin regulators (KRAB and DNMT3A) or by recruitment of endogenous chromatin regulators by other factors as represented by *α*, *β* and *β*_0_. The green arrows represent H3K9me3 decay while the blue dashed arrows represent the erasure of DNA methylation through TET1. U represents unmodified nucleosomes, x represents nucleosomes with H3K9me3, y represents nucleosomes with DNA methylation, and z represents nucleosomes with both marks. Positive feedback marked by *α* represents H3K9me3 autocatalysis. Arrows marked by *β* represent the recruitment of DNA methylation by H3K9me3. Arrows marked by *β*_0_ represent the recruitment of H3K9me3 by DNA methylation. Only some of these recruitment arrows are shown to avoid clutter. **B.** Trajectories of the fraction of H3K9me3 methylation (grey) and DNA methylation (black) after temporary recruitment of DNMT3A followed by temporary recruitment of TET1. **C.** Trajectories of the fraction of H3K9me3 methylation (grey) and DNA methylation (black) after temporary recruitment of KRAB followed by temporary recruitment of TET1. Red shade indicates the timespan of DNMT3A or KRAB recruitment whereas the blue shade indicates the timespan of TET1 recruitment. Computational simulations were performed with the ODE model in Supplementary Text. **D.** In the absence of DNMT3A or KRAB and presence of high TET1 levels, the circuit in panel A reduces to a two-state system devoid of DNA methylation. Here, U represents the fraction of unmodified nucleosomes and x represents the fraction of H3K9me3-modifed nucleosomes. The ODE represents the rate of change of the fraction of modified nucleosomes, where *α* represents the strength of H3K9me3 autocatalysis and *δ* represents its decay constant. **E.** The steady states lie at the intersection of the production rate (black) and decay rate (green) lines. Solid circles represent stable steady states while open circles represent unstable steady states.

Positive feedback thus emerges around the state z because DNA methylation and H3K9me3 recruit each other (arrows marked by *β* and *β*_0_). This positive feedback is lacking in CHO-K1 cells (*β* = 0, diagram in Fig. S7A-B, left), in which H3K9me3 does not recruit DNA methylation ^65^. Moreover, H3K9me3 is autocatalytic as it recruits its own writers, what is known as heterochromatin spreading ^93–95^ (feedback loop marked by *α* in Fig. 3A). Finally, H3K9me3 decays passively through dilution and actively by the action of histone demethylases (green arrows in Fig. 3A, marked by *δ*). In contrast, DNA methylation is stable due to the maintenance action of DNMT1 when TET1 eraser enzyme is absent ^96,97^. Therefore, the decay rate of DNA methylation is negligible when no TET1 is recruited but increases proportionally with the amount of TET1 present (blue arrows in Fig. 3A). The full set of reactions is reported in Supplementary Text.

### Recapitulating silencing dynamics

According to the model, starting from an active gene state that lacks DNA methylation and H3K9me3, transient recruitment of DNMT3A will lead to the permanent establishment of DNA methylation, which will, in turn, lead to stable establishment of H3K9me3 (Fig. 3B and S7B). Transient recruitment of KRAB will lead to H3K9me3, which, in turn, will recruit DNA methylation in hiPSCs (*β* ≠ 0) but not in CHO-K1 cells (*β* = 0). If this recruitment is sufficiently fast (*β* sufficiently large) compared to the rate at which H3K9me3 decays, then DNA methylation will accumulate in the gene and lead to stable silencing (Fig. 3C, before time T). By contrast, in CHO-K1 cells, transient recruitment of KRAB will lead to transient establishment of H3K9me3, no DNA methylation, and hence to only transient silencing (Fig. S7A). Consequently, in hiPSCs (*β* ≠ 0) we observe permanent silencing in both cases, consistent with the experimental data pertaining to the hEF1a promoter (Fig. 2D-E).

Nevertheless, if the rate of *de novo* DNA methylation is exceedingly low such that it is mostly established through the recruitment of DNMT3A by H3K9me3, then it is possible that DNA methylation can only be established when both KRAB and DNMT3A are concurrently recruited to the promoter, but not by DNMT3A alone (Fig. S8, middle and bottom plots). Similarly, the effective recruitment of DNA methylation by H3K9me3 hinges on a sufficiently large rate constant *β*, which increases with the local availability of free DNMT3A and H3K9me3-binding proteins that recruit DNMT3A. For low values of the rate constant, H3K9me3 alone will not recruit sufficient DNMT3A to stabilize itself, resulting in the decay of H3K9me3 and transient silencing (Fig. S8, top plot). Exogenous recruitment of DNMT3A (via transient transfection) in the presence of H3K9me3 can increase this rate and enable the establishment of DNA methylation, leading to durable silencing. This rate-dependent behavior is in agreement with the experimental data pertaining to the CAG promoter in hiPSCs (Fig. S3B).

### Predicting reactivation dynamics

In the presence of high levels of TET1, the chromatin modification circuit of Fig. 3A will be devoid of DNA methylation (DNAme), such that the system reduces to the two-state circuit of Fig. 3D. In this circuit, unmodified nucleosomes (U) become modified to H3K9me3 (x) through positive feedback (*α*), and modified nucleosomes become unmodified through decay (*δ*). The steady state values of the fraction of H3K9me3 (*x*) are obtained when the production and decay terms are equal: *α* · *x* · (1 − *x*) = *δ* · *x* (Fig. 3E, and Supplementary Text). When *α* ≫ *δ*, that is, the autocatalysis overpowers decay, two steady states appear, but only the non-zero one corresponding to saturating H3K9me3 located at *x* = (*α* − *δ*)*/α* ≈ 1 is stable (the silenced state). Conversely, when *α* ≪ *δ*, meaning decay overpowers autocatalysis, the system has only one stable steady state at *x* ≈ 0 completely lacking H3K9me3 (the active state). Therefore, only in the latter case can we expect gene reactivation after transient TET1 recruitment since removal of DNA methylation will also lead to H3K9me3 decay.

Data in CHO-K1 cells indicated that H3K9me3 could not sustain itself in the absence of DNA methylation (S1E) ^65^, which was recapitulated in hiPSCs when using the CAG promoter (Fig. S6). These data thus suggest that H3K9me3 decay overpowers its autocatalysis (*α* ≪ *δ*). In this case, the model predicts durable gene reactivation, assuming that DNA de-methylation by TET1 is highly efficient (Fig. 3B-C, after time T and *α* ≪ *δ*, and Fig. S8, bottom). If the efficacy of DNA demethylation by TET1 is low, residual DNA methylation will reinforce H3K9me3 (through *β*_0_), which will reciprocally reinforce DNA methylation (through *β*). The system could then restore DNA methylation and H3K9me3 levels after removal of TET1 if this positive feedback is sufficiently strong, thereby silencing the locus again. In this case, TET1 alone would lead to transient reactivation, possibly requiring a combination of TET1 and H3K9me3 erasers to achieve durable reactivation. In summary, for both marks to be permanently erased after transient recruitment of TET1, it is necessary that demethylation is highly efficient and that the positive feedback loop between DNA methylation and H3K9me3 is not strong enough to build up the repressive marks starting from any residual DNA methylation level (Fig. S9).

### Reversing induced epigenetic silencing in hiPSCs

The model predictions suggested that similar reactivation to that observed in CHO-K1 cells could occur in hiPSCs if DNA demethylation by TET1 is highly efficient and H3K9me3 autocatalysis is overpowered by its decay. We therefore attempted reactivating the reporter system using rTetR-TET1 in hiPSCs previously silenced with DNMT3A or KRAB. To probe reactivation dynamics, we co-transfected cells with rTetR-TET1 and EYFP transfection marker, isolated transfected cells (EYFP+) at day 2 by FACS and periodically measured reporter gene expression by flow cytometry (Fig. 4A). As predicted by the model, DNMT3A-silenced iPSCs transfected with rTetR-TET1 and cultured in dox for 6 days displayed strong reactivation of gene expression 7 days post-transfection, which remained stable for the duration of the experiment (Fig. 4B). Transfection marker (EYFP) levels indicate that the chromatin regulator was completely diluted out by day 7, implying that expression levels were stable in the absence of the regulator. Interestingly, when we repeated the experiment using KRAB-silenced hiPSCs, we similarly observed efficient and stable reactivation of transgene expression up to 15 days post-transfection (Fig. 4C). Chromatin analysis of DNA methylation and H3K9me3 revealed that both marks were depleted after rTetR-TET1 recruitment in both cases (Fig. 4D), in line with model predictions (Fig. 3B-C). Methyl CpG sequencing at the operator-promoter regions also showed a return to baseline methylation levels after TET1 treatment of KRAB-silenced hiPSCs, demonstrating the efficacy of TET1 in reactivating KRAB-silenced locus (Fig. 4E and S10A). These data corroborate the hypothesis that H3K9me3 autocatalysis is not strong enough compared to the mark’s decay rate and hence that it cannot sustain itself in the absence of DNA methylation. As a consequence, it is possible to reactivate silenced transgene loci by only removing DNA methylation even when there are repressive histone marks without necessitating direct H3K9me3 erasure.

**Fig. 4.**
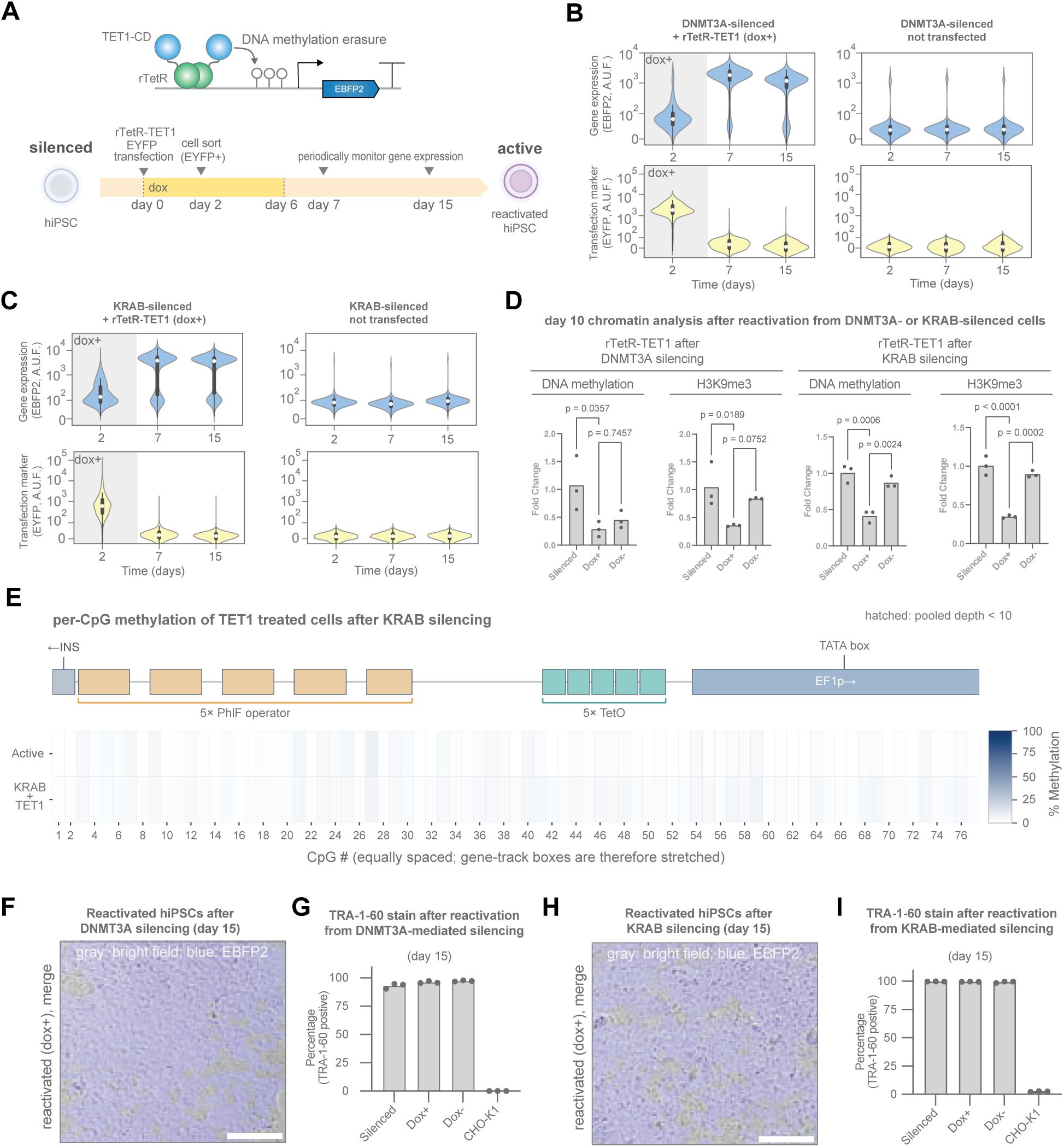
Targeted chromatin editing with an engineered TET1-based chromatin regulator efficiently reverses induced silencing in hiPSCs. **A.** Schematic of the experimental setup for hiPSC reactivation by transient rTetR-TET1 transfection. **B-C.** Violin plots of flow cytometry measurements after rTetR-TET1 transfection of hiPSCs silenced with **(B)** rTetR-DNMT3A or **(C)** PhlF-KRAB. Data are plotted from three independent replicates. The white dot represents the median, the thick box represents the interquartile range (IQR) and the thin gray line represents 1.5x the IQR. **D.** Quantitative PCR analysis (qPCR) of chromatin after methylated DNA immunoprecipitation (MeDIP) or CUT&RUN targeting H3K9me3 (see Methods). Bars represent mean values of three independent replicates. Statistics were performed with an ordinary one-way ANOVA with Dunnet correction for multiple comparisons. **E.** Heatmap of targeted TAPS+ sequencing of the reporter transgene promoter. Methylation level of each CpG residue within the operator and promoter regions are shown for active and TET1-reactivated cells after KRAB-silencing. Data reported are average from three independent replicates. **F-G.** Pluripotency assessment by **(F)** morphology and **(G)** cell surface marker staining (TRA-1-60) of reactivated cells from DNMT3A-silenced state. **H-I.** Pluripotency assessment by **(H)** morphology and **I)** cell surface marker staining (TRA-1-60) of reactivated cells from KRAB-silenced state. Scale bars are 150 *µ*m.

To confirm that pluripotency was unaffected by epigenome editing with TET1 overexpression, we performed live cell microscopy to assess colony morphology and transgene expression in addition to quantification of cell surface marker expression for undifferentiated pluripotent stem cells (TRA-1-60) ^98^ by flow cytometry (Fig. 4F-I). Cells reactivated following DNMT3A-silencing displayed colonies with smooth edges and high EBFP2 fluorescence (Fig. 4F and S10B) while maintaining >90% of the population expressing the TRA-1-60 marker of undifferentiated pluripotent stem cells (Fig. 4G). Similarly, KRAB-silenced iPSCs reactivated with TET1 formed colonies with smooth edges and devoid of spontaneously differentiated cells while expressing EBFP2 (Fig. 4H and S10C), and the vast majority of the population stained positive for the TRA-1-60 surface marker (Fig. 4I). Unlike dCas9-VPR, transient recruitment of TetR-TET1 to the silenced CAG-driven reporter also reactivated gene expression, suggesting that the reactivation function of TET1 generalizes across promoters (Fig. S10D). Together, these results indicate that our targeted chromatin editing does not negatively impact stem cells while still managing to transcriptionally modulate the transgene locus for durable reactivation in hiPSCs.

### Reversing endogenous epigenetic silencing in hiPSCs

Motivated by the result that transgene reactivation was possible from silenced states with both DNA methylation and H3K9me3, we asked whether we could reverse endogenous epigenetic transgene silencing. To this end, we first isolated hiPSCs in the active state (EBFP2+) by FACS and cultured them under standard conditions for 12 days, anticipating the emergence of an endogenously-silenced population (EBFP2-) that we subsequently isolated by FACS (Fig. 5A, bottom panel red gate). We then repeated the experiment outlined in Fig. 4A but starting with the EBFP2-sorted population. The cells transfected with rTetR-TET1 (EYFP+) and cultured in dox for 6 days showed efficient reporter gene reactivation at day 7 that persisted until day 15 (Fig. 5B). A small increase in the fraction of silenced cells is noticeable between days 7 and 15, which is likely due to endogenous silencing since they are lacking rTetR-TET1 based on EYFP expression. Chromatin analyses of the promoter region revealed that rTetR-TET1 recruitment (dox+) led to a significant decrease in the levels of both DNA methylation and H3K9me3 when compared to the silenced cells (Fig. 5C). Per-CpG methylation analysis of endogenously-silenced hiPSCs showed the highest percent-methylation (70-90%) of all samples, which we were able to efficiently reverse by transient TET1 recruitment (Fig. 5D and S11A). We also tested other activator domains to compare their ability to reactivate gene expression in hiPSCs (Fig. S11B). For the rTetR-P300 fusion protein, we could not detect reporter expression after 2 or 8 days post transfection. Next, the combination of rTetR-PRDM9 and rTetR-DOT1L initially displayed slight EBFP2 fluroescence at day 2, but the activation was not stable and decreased to the initial silenced level by day 8, suggesting that recruitment of a histone regulator is not conducive to reactivation at a heavily methylated locus, however, we have not measured expression levels of these fusion proteins or assayed whether their cognate mark has been deposited at / near the promoter region. We observed similar behavior when targeting dCas9-VPR to all TetO sites, with an initial burst in fluorescence that faded with time. In fact, the only durable activation we observed was when transfecting rTetR-TET1. These cells displayed a smaller increase in fluorescence at day 2 when compared to DOT1L/PRDM9 or VPR targeting, but showed strong fluorescence signal at day 8. To determine whether the time spent in silenced state affected reactivation efficiency, we cultivated endogenously-silenced hiPSCs for >4 weeks (Fig. S11C), then subjected them to transient transfection with TET1 or catalytically inactive TET1 (deadTET1) and measured reporter gene expression. We observed similar results, where TET1 targeting led to strong reactivation while mock transfection or inactive TET1 did not show signs of reactivation (Fig. S11D). These data demonstrate that transgene loci endogenously silenced by DNA methylation and H3K9me3 can be stably reactivated by removing DNA methylation only, which then leads to an automatic loss of the H3K9me3 mark. Reactivated cells maintained their undifferentiated pluripotent state, as is shown by microscopy analysis of colony morphology (Fig. 5E and S12A) and expression of cell surface marker TRA-1-60 (Fig. 5F).

**Fig. 5.**
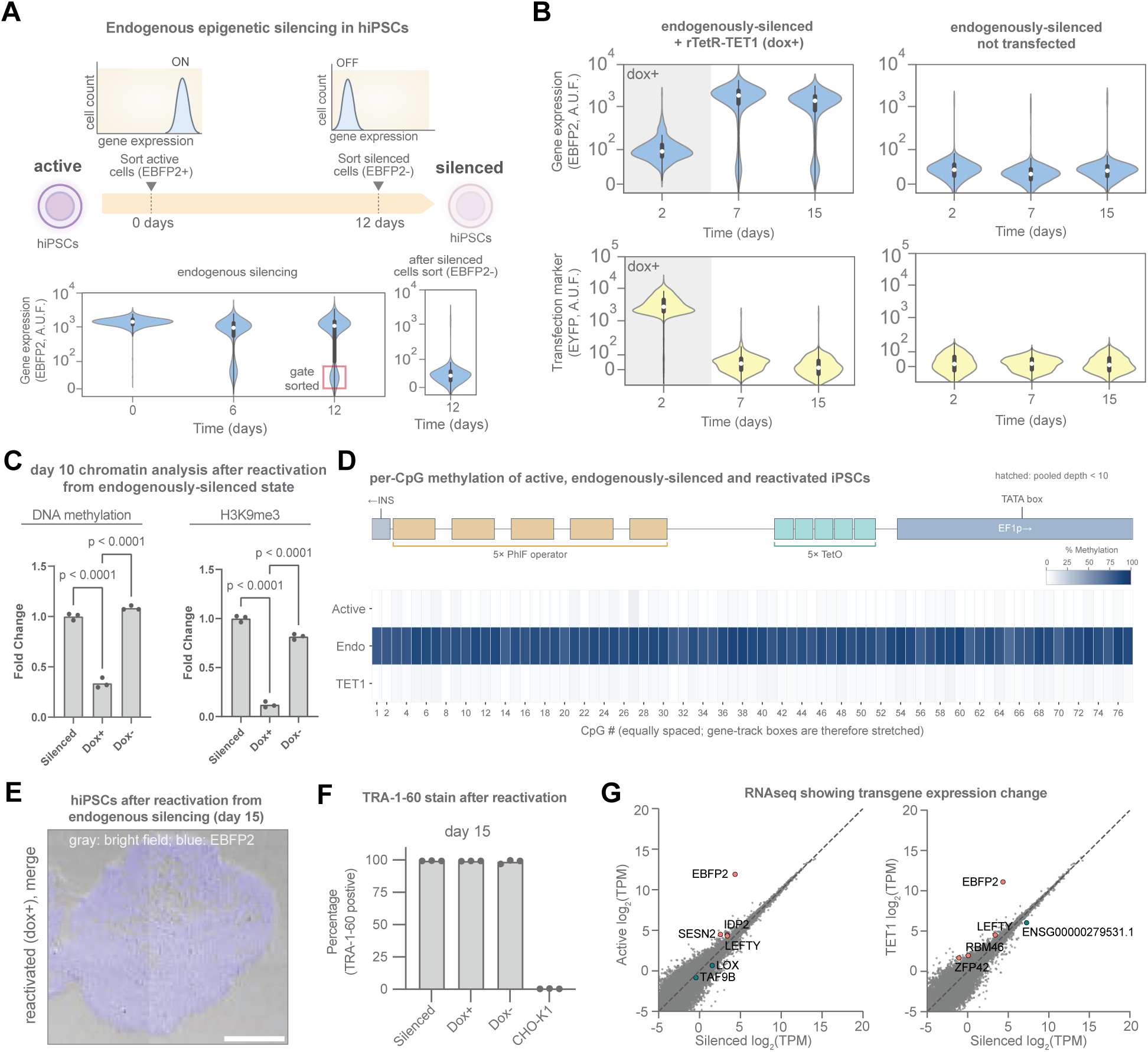
Endogenous transgene silencing in hiPSCs is efficiently reversed by targeted TET1-mediated demethylation. **A.** Top: Schematic of the experimental setup to obtain endogenously silenced hiPSCs. Bottom: Violin plots of flow cytometry measurements tracking reporter fluorescence. At day 12, non-expressing cells (EBFP2-) were sorted (red gate), expanded and used as starting point for reactivation experiments. **B.** Violin plots of flow cytometry measurements after rTetR-TET1 transfection of endogenously-silenced hiPSCs. Error bars are s.d. of the mean from three independent replicates. **C.** Quantitative PCR analysis (qPCR) of chromatin after methylated DNA immunoprecipitation (MeDIP) or CUT&RUN targeting H3K9me3. Bars represent mean values of three independent replicates. Statistics were performed with an ordinary one-way ANOVA with Dunnet correction for multiple comparisons. **D.** Heatmap of targeted TAPS+ sequencing of the reporter transgene promoter. Methylation level of each CpG residue within the operator and promoter regions are shown for active, endogenously-silenced and TET1-reactivated cells. Data reported are average from three independent replicates. **E-F.** Pluripotency assessment by **(E)** morphology (scale bar is 300 *µ*m) and **(F)** cell surface marker (TRA-1-60) staining of reactivated cells from the endogenously-silenced state. Data are plotted as mean from three independent replicates. **G.** Normalized transcript counts for cells in the active state, endogenously-silenced cells, and reactivated with TET1. TPM: Transcripts per million. Red is upregulated, green is downregulated. For all violin plots, the white dot represents the median, the thick box represents the interquartile range (IQR) and the thin gray line represents 1.5x the IQR.

We further corroborated the data by performing bulk RNAseq on active cells as well as endogenously silenced cells before and after reactivation with TET1 to confirm specific targeting and exclude any off-target effects (Fig. 5G and S12B-C). The data clearly show downregulation of EBFP2 between active and silenced cells and subsequent upregulation between silenced and reactivated (TET1) cells. Importantly, when comparing active cells to reactivated cells (Fig. S12C), we found no difference in gene expression profiles between the two samples, indicating that reactivation did not result in undesired transcriptional changes and that the re-activated cells returned to their original state before endogenous silencing. We did observed a slight non-specific transcriptional up-regulation of a few endogenous genes between silenced and reactivated (TET1) cells. In particular, ZFP42 or REX1 (on chromosome 4q35), RBM46 (on chromosome 4q32), and LEFTY1 (on chromosome 1q42) were upregulated in reactivated cells relative to silenced cells. The ZFP42 transcription factor showing highest expression fold change is broadly associated with pluripotency maintenance, consistent with a transient activation of pluripotency genes ^99^, while LEFTY1, a secreted inhibitor of SMAD signalling ^100^, borders the expression fold change limit and is associated with early developmental gene modules. RBM46 instead displays an intermediate expression fold change with respect to the other two genes, and is a testis- and germline-enriched RNA-binding protein localizing to the cytoplasm ^101^. Despite these minor differences, it is noteworthy that the relative endogenous gene expression changes between silenced and reactivated cells are thwarted by the significant transcriptional modulation elicited on the targeted EBFP2 reporter transgene. Overall, the results suggest that the our system for TET1-mediated transgene reactivation is highly specific to the targeted locus and leads to little off-target effects while restoring transgene expression to its original levels.

### Lentiviral integrations display similar dynamics and can be rescued from silencing

To assess whether our results are generalizable to different chromatin contexts, we sought to integrate the reporter transgene via lentiviral transduction to obtain a population of expressing cells with the reporter found in random locations within the genome. To this end, we built the reporter construct flanked by LTRs and swapped the terminator with a WPRE, transduced wild-type CHO-K1 and hiPS cells, and generated EBFP2+ polyclonal cell lines by FACS (Fig. 6A). We transiently transfected the CHO-K1 cells with rTetR-DNMT3A or PhlF-KRAB along with a transfection marker (EYFP), sorted transfected cells by FACS and monitored reporter gene expression over time. The silencing dynamics mirror those obtained with site-specific reporter integration, where DNMT3A-treated cells displayed durable silencing whereas KRAB-treated cells showed only transient silencing that inversely correlated with KRAB expression (Fig. S13A). We then subjected the DNMT3A-silenced CHO-K1 cells to transfection with TetR-TET1 and followed reporter gene expression levels after sorting. We could observe strong reactivation of the reporter at day 6 that persisted until day 12 (Fig. 6B), while control transfections with plasmid backbone (mock) or catalytically inactive TET1 (deadTET1) led to no detectable gene expression.

**Fig. 6.**
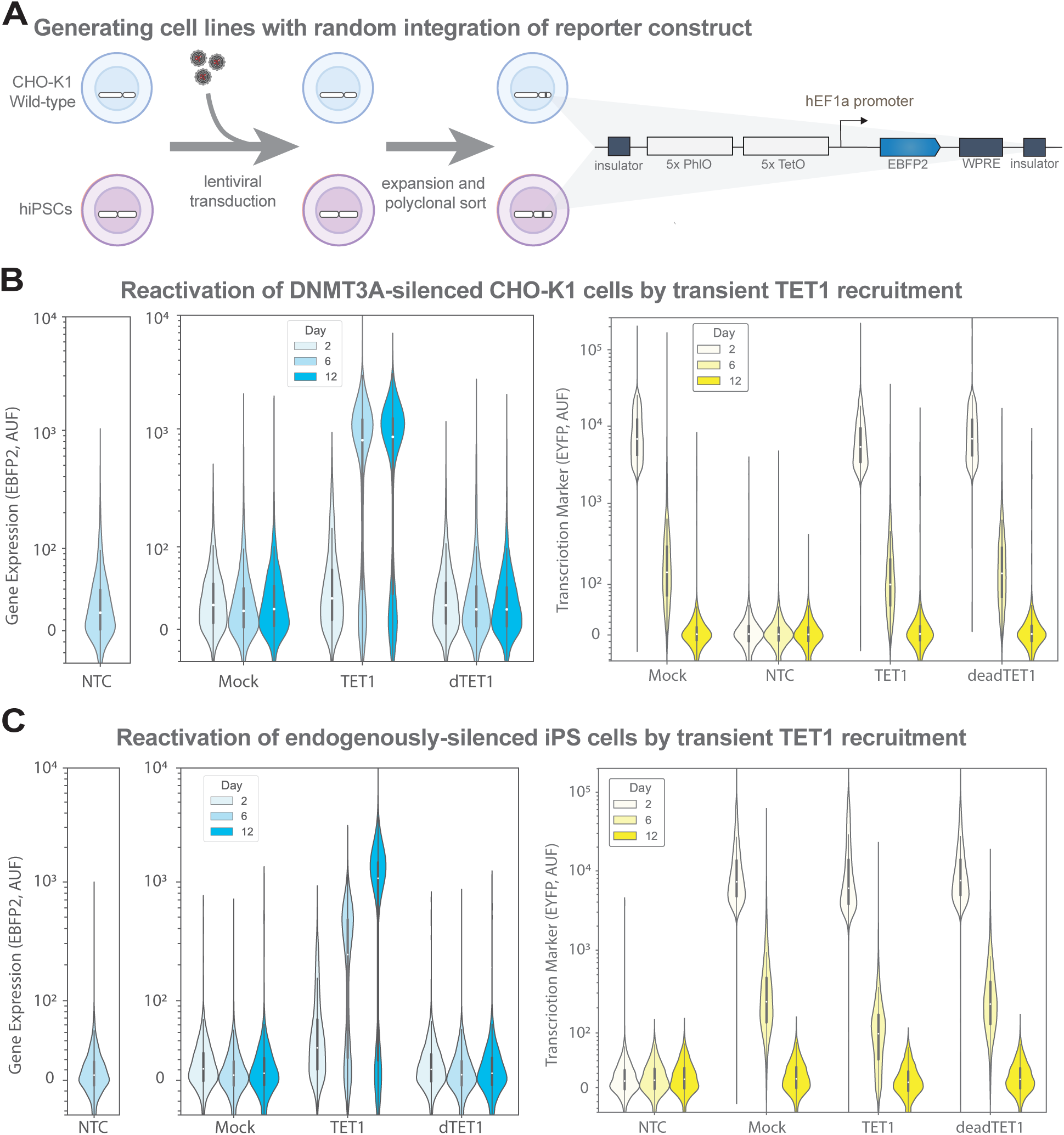
Targeted TET1 recruitment reactivates lentivirally-integrated silenced reporter in both CHO-K1 and hiPS cells. **A.** Schematic showing the generation of reporter cell lines via lentiviral transduction. The construct is equivalent except for the terminator element which was swapped for WPRE to enhance lentivirus generation and expression. **B.** Violin plots showing TET1-mediated reporter reactivation (left) and transfection marker levels (right) of DNMT3A-silenced CHO-K1 cells. **C.** Violin plots showing TET1-mediated reporter reactivation (left) and trasnfection marker levels (right) of endogenously-silenced hiPS cells. For all violin plots, Mock transfection is an empty plasmid backbone. deadTET1/dTET1 is catalytically inactivated TET1. The white dot represents the median, the thick box represents the interquartile range (IQR) and the thin gray line represents 1.5x the IQR.

Next, we moved to probe the dynamics of the lentivirally-integrated reporter in hiPSCs. Repression by transient transfection of either DNMT3A or KRAB resulted in durable silencing that persisted until day 12, long after the regulator was diluted (Fig. S13B), recapitulating the results obtained with the reporter in the AAVS1 locus (Fig. 2D-E). Thus, the rules governing the silencing dynamics of the reporter transgene are agnostic to the chromosomal locus or integration strategy employed. Differently from the previous results though, reactivation efficiency was not comparable to that observed in the AAVS1 locus starting from DNMT3A- or KRAB-silenced cells, achieving 30% reactivation for both conditions. We attribute this discrepancy between the two cases to an increased toxicity of TET1 in transduced cells that had previously been repressed by transient transfection, such that when sorting cells by FACS 2 days after transfection, the population distributions of EYFP+ cells were skewed to towards lower levels, which in turn suggests that TET1 levels were accordingly low (Fig. S13C, compare EYFP values at day 2).

Finally, we aimed to evaluate reactivation efficiency of the lentiviral reporter after endogenous silencing. We therefore cultured the active cells until a silenced population emerged and subsequently sorted and expanded that population. When we targeted TET1 to these cells by transient transfection, we could detect reporter gene expression from the lentivirally-integrated transgene that persisted until day 12, indicating durable, efficient reactivation and reversal of endogenous silencing (Fig. 6C). This observation reinforces the hypothesis that in the previous experiment with DNMT3A and KRAB edited cells inefficient reactivation was likely due to toxicity from the second round of transfections with TET1. Based on these results, the silencing and reactivation dynamics of the reporter transgene appear to hold independent of the chromosomal location or chromatin state. Thus, a model emerges where durable transgene silencing can be achieved with DNMT3A targeting in CHO-K1 cells that can be reversed by targeted TET1 activity independent of the promoter used or chromatin context (Fig. 7A), while durable silencing in iPSCs is promoter dependent (hEF1a promoter requires DNMT3A or KRAB while CAG promoter necessitates both), although both silenced states can be reactivated with targeted TET1 treatment only (Fig. 7B).

**Fig. 7.**
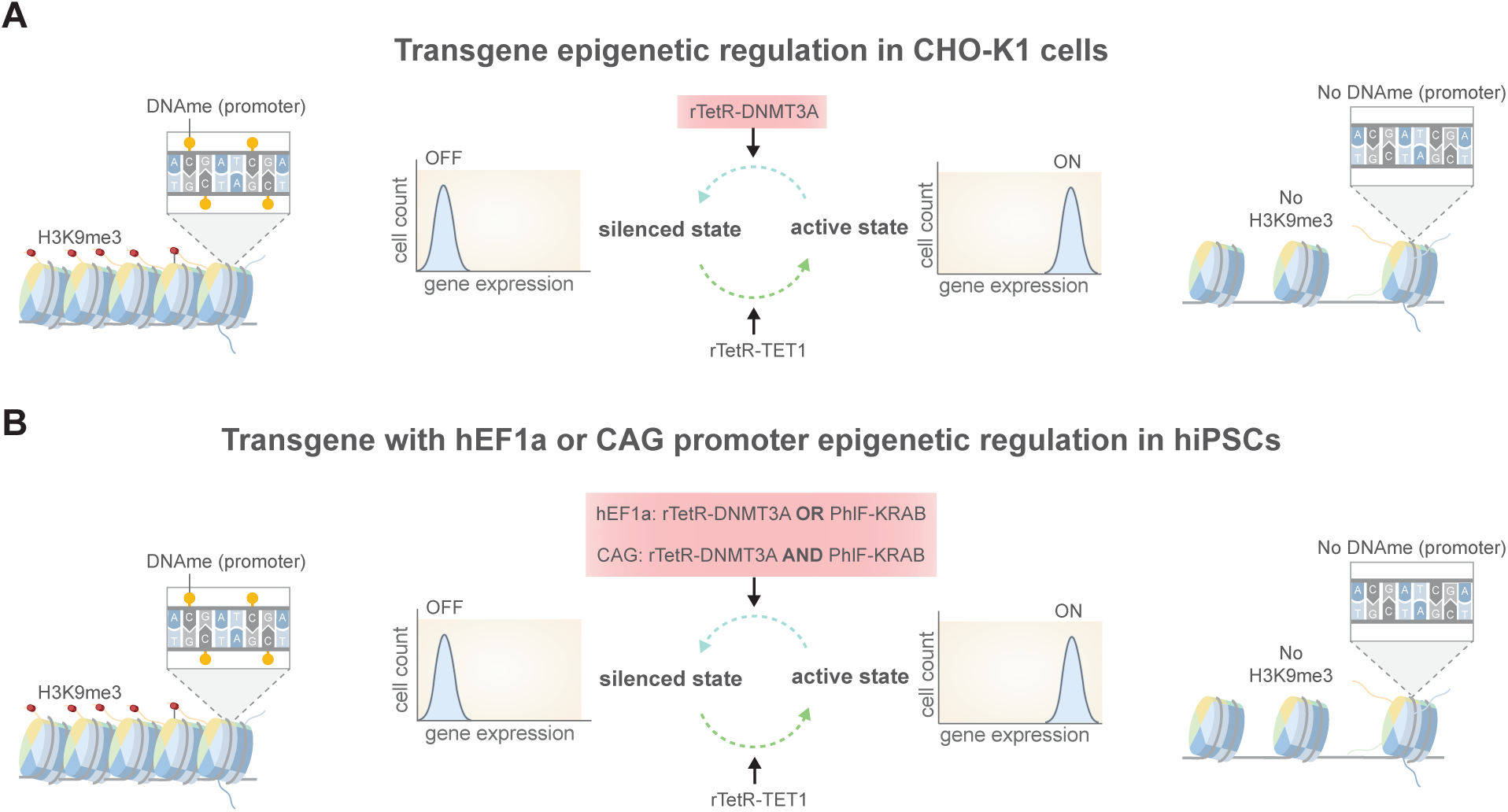
Schematic showing emergent model for context-dependence of transgene transcriptional control by chromatin editing. **A.** Model of reversible transgene epigenetic editing in CHO-K1 cells cells. **B.** Model of reversible transgene epigenetic editing in hiPSCs with hEF1 or CAG promoter.

## Discussion

While epigenome engineering has made great strides in understanding chromatin dynamics, a systematic investigation within a controlled genetic context of the molecular mechanisms at play in transgene silencing and reactivation had been lacking. Here, we developed a reporter system in two different cell types that differ in the locus of integration, but are otherwise identical, and used targeted chromatin editing to investigate how chromatin marks interact during epigenetic silencing and reactivation (Fig. 1, 2 and 4). We demonstrated that transient expression of rTetR-DNMT3A durably silences the transgene in both cell types and that it leads to an increase in both DNA methylation and H3K9me3 at the promoter region (Fig. 1C-D, 2D, F). In contrast, transient expression of PhlF-KRAB led to stable silencing in hiPSCs but only to transient repression in CHO-K1 cells (Fig. 2E and S1E, respectively). Chromatin analysis revealed an enrichment of H3K9me3 in both cell types and a significant increase in DNA methylation in hiPSCs but not in CHO-K1 cells (Fig. 2F and ref ^65^). We further analyzed the consequence of direct H3K9me3 deposition by targeting the G9A SET domain, which in principle excludes the possibility of co-recruitment of DNMT3s reported for the KRAB domain ^102–104^. Transient transfection with this chromatin regulator again led to both H3K9me3 and DNA methylation deposition (Fig.S5B-D), although to a lesser extent when compared with KRAB recruitment, indicating that the deposition of H3K9me3 itself triggers recruitment of DNA methylation. Taken together, these data indicate that a positive feedback loop between H3K9me3 and DNA methylation exists in hiPSCs but is absent in CHO-K1 cells.

We used a mathematical model of chromatin modifications to recapitulate silencing dynamics and make predictions on how reactivation dynamics are shaped by the interaction among H3K9me3, DNA methylation and network parameters (Fig. 3). The model is adopted from our earlier work ^89^, but only includes the dynamics of the repressive chromatin marks. The model indicates that the positive feedback loop between DNA methylation and H3K9me3 leads to durable silencing after deposition of H3K9me3 even when H3K9me3 turnover overpowers the rate of autocatalysis such that the mark is unable to sustain itself. In this situation, the model predicts that it is possible to reactivate gene expression by targeted DNA demethylation despite having repressive histone marks (H3K9me3) at the locus (Fig. 3B-C; top panels) because repression is maintained by DNA methylation. The experiments with transient expression of rTetR-TET1 in silenced cells confirmed that we can rescue transgene expression from a silenced gene state by only removing DNA methylation and that both DNA methylation an H3K9me3 are diminished after treatment (Fig. 1C-D, S1C-D, 4B-E, 5B-D, S10A, S11A,D). Nevertheless, the model indicates that even with weak H3K9me3 autocatalysis, a sufficiently strong positive feedback between DNA methylation and H3K9me3 may prevent stable reactivation by targeted TET1, since residual levels of DNA methylation will recruit H3K9me3, which in turn will trigger recruitment of more DNA methylation, ultimately leading to their accumulation and subsequent gene silencing (Fig. S9). The strength of this feedback is context-dependent and linked to the availability of endogenous DNMT3s and H3K9me3 readers such as HP1 (*β*) and of histone methyltransferases and methyl-CpG binding proteins that read DNA methylation (*β*_0_). Related models have appeared in the literature that include both H3K9me3 and DNA methylation, however the positive feedback between these two marks was not captured ^105^.

We observed that the reporter system in CHO-K1 cells transiently transfected with PhlF-KRAB display only transient silencing with expression levels returning to normal after PhlF-KRAB dilution (Fig. S1E and ref ^65^), concordant with other reports using CHO-K1 cells and the KRAB domain ^46^. Targeting of the KRAB domain was also reported to lead to transient repression ^45,49,69^, thus many studies have opted for the co-recruitment of DNMT3A/L with KRAB to exhibit stable silencing ^58,68,70^. These observations suggest that while H3K9me3 recruitment by KRAB represses gene expression, silencing that persist after KRAB removal may be achieved by also recruiting DNA methylation in these genomic contexts. Contrarily, when we repeated the KRAB-silencing experiment in hiPSCs with the hEF1a-driven reporter, we observed stable silencing (Fig. 2E and S13B). This discordant behavior between CHO-K1 and hiPS cells might be attributed to differing chromatin regulatory networks governing DNA methylation and H3K9me3. Specifically, H3K9me3 leads to the recruitment of DNA methylation in hiPSCs (Fig. 2F-G) but not in CHO-K1 cells ^46,65^, indicating that the permanence of H3K9me3 in hiPSCs may be due to the positive feedback between DNA methylation and H3K9me3 (Fig. 3A), which is instead lacking in CHO-K1 cells ^65^ (Fig. S7). Intriguingly, a study performed in mouse embryonic stem cells (mESCs) showed accumulation of DNA methylation after targeting of KRAB-MeCP2 fusion construct, but also observed stable silencing in DNMT3A/3B double knockout mESCs after transient recruitment of KRAB-MeCP2, independent of DNA methylation ^106^. In our model, absence of DNMT3s leads to *β* = 0, and therefore stable silencing can be observed only if the autocatalysis of H3K9me3 overpowers its decay (Fig. 3C,D,E). It is therefore possible that the knockout cell lines had higher levels of H3K9 methyltransferases and / or of HP1 proteins, both necessary for H3K9me3 autocatalysis. A different study that aimed to assess genome-wide transposon silencing upon global DNA demethylation found that the cell ensures genomic stability by maintaining transposons silenced with repressive histone marks even in the absence of DNA methylation. These observations suggest that, while DNA methylation plays a critical role in maintenance of repressed state when possible, cells are nonetheless capable of durable silencing with H3K9me3 or H3K27me3 in cases where DNA methylation is abolished ^107^.

The context-dependence of the network of chromatin modifications can be attributed to the differential presence of molecular players enabling the interactions, which in turn may be conditioned to the specific cell type, genomic integration site, and / or promoter. We confirmed that the dynamics observed are not conditioned to the specific integration site by performing experiments on the same reporter randomly integrated in a polyclonal population. The results resembled those obtained with site-specific integration (Fig. 6, S13), suggesting that the presence of the positive feedback between DNA methylation and H3K9me3 in our system is not a peculiarity of the genomic integration site. Instead, differences in cell type may be a likely cause. For instance, the protein UHRF1, consisting of Tandem-Tudor domain (TTD), plant homeodomain (PHD) and SET and RING associated domain (SRA), can bind H3K9me2/3, H3K4me0/1 and hemimethylated DNA in addition to recruiting maintenance methylase DNMT1 and *de novo* methylases DNMT3A/B ^108–112^. While this protein has been observed experimentally in humans (*Homo sapiens*), it is only predicted to be present in hamsters (*Cricetulus griseus*) (see UniProt entries Q96T88 and A0A8C2LY95, respectively), which might explain why H3K9me3 leads to DNA methylation in hiPSCs but not in CHO-K1 cells. However, the study showing reversibility of KRAB-induced silencing of CAG driven reporter in human iPSCs but stable silencing in hiPSC-derived cardiomyocytes ^113^, similar to what we observed with CAG reporter silencing with KRAB (Fig. S6), suggests that these parameters may also depend on the promoter used and / or cell differentiation state.

Our data indicate that a variety of commonly used mammalian promoters, including CAG, murine PGK, and human UbC promoters, can be similarly silenced and re-activated in CHO-K1 cells, despite differences in expression levels and sequence compositions (Fig. S3). However, we found that the silencing and reactivation dynamics of the CAG promoter were different from that of the hEF1a promoter in hiPSCs (Fig. 2E and S6). Specifically, KRAB-based chromatin editing of CAG-driven expression led to transient silencing, as in CHO-K1 cells and as reported elsewhere ^113^, while DNMT3A-based editing did not exert any effect. However, combining both chromatin regulators resulted in durable silencing (Fig. S6), which could be reversed by TET1-mediated reactivation (Fig. S10D), similarly to what we observed for the hEF1a promoter in hiPSCs and CHO-K1 cells.

The model recapitulates these differences when DNA methylation recruitment by H3K9me3 necessitates rTetR-DNMT3A at the promoter (Fig. S8). The strength of this recruitment (*β*) is proportional to the availability of both DNMT3A at the promoter and H3K9me3-binding proteins that recruit DNMT3A. If there is an abundance of the latter but not of the former around the promoter, recruitment of H3K9me3 alone is insufficient to recruit endogenous DNMT3A, resulting in transient repression after KRAB editing. When we co-express rTetR-DNMT3A in the presence of H3K9me3, this can more effectively recruit DNMT3A to establish stable DNA methylation and lead to durable repression. Instead, without H3K9me3, rTetR-DNMT3A recruitment to the locus may be insufficient to establish stable DNA methylation, leading to no effect on the reporter gene’s expression.

A potential mechanism for how H3K9me3 may enable the activity of DNMT3s is via loss of active chromatin marks such as H3K4me3. The structure of DNMT3 family comprise an ATRX-DNMT3-DNMT3L (ADD) domain that allosetrically inhibits the catalytic domain, which can be relieved upon binding to H3K4me0^114^, and in this way directs endogenous DNMT3s to inactive regions of the genome. Thus, since the KRAB domain associates with KAP1/TRIM28, which in turn recruits SETDB1, a H3K9me writer, Mi2*α*, a component of the nucleosome remodeling and histone deacetylation (NuRD) complex, and HP1^102^, the concerted action of these repressor complexes could potentially lead to the loss of activating histone marks (increased H3K4me3 demethylation to H3K4me0) and subsequent DNMT3 recruitment and DNA methylation deposition. Interestingly, H3K14ac has been shown to protect methylated H3K4 demethylation from lysine specific demethylase 1 (LSD1/KDM1A) ^115,116^, so H3K14 deacetylation by the NuRD complex may allow H3K4 demethylation by LSD1^117^. Nevertheless, further studies are needed to elucidate the exact mechanism of interplay between the different chromatin marks and DNA methylation.

Prior work investigated the reactivation of endogenous genes that were either hypermethylated or unmethylated with PRDM9 or VP64, concluding that reactivation cannot be sustained for an extended period of time without first removing DNA methylation at the locus, which they showed with a nucleoside analog ^79^. Our data corroborates this conclusion for transgenes, since we were unable to durably reactivate the silenced reporter with the other chromatin regulators or transcriptional activators tested in either CHO-K1 (Fig. 1E and S2C) or hiPSCs (Fig. S11C). Other studies have shown gene reactivation by targeted DNA demethylation in both endogenous and integrated genes in a variety of cell lines, suggesting that reactivation was possible for genes silenced by DNA methylation ^45,50,57,64,68,73,118,119^, although none demonstrated stable reactivation of transgene integrations in hiPSCs. Notably, we have shown that TET1 recruitment to a KRAB-silenced transgene locus in iPSCs also leads to stable gene reactivation (Fig. 4C-E), which we extended to reliably reactivate the endogenously silenced reporter (Fig. 5B-D). We did observe a small increase in the fraction of silenced cells between days 7 and 15, indicating that while the reporter was reactivated, it nonetheless remained susceptible to endogenous silencing. These results are of relevance for engineering cell fates since transgenes have been implicated as targets for the human silencing hub (HUSH) complex that ultimately leads to H3K9me3 and subsequent DNA methylation deposition ^17,19,120,121^. The complex typically serves to maintain genome integrity by repressing retroviral elements, which it recognizes via their intronless mRNAs. Consequently, synthetic genetic circuits that express proteins from cDNA also become targets of the HUSH complex. Thus, for cell engineering of iPSCs, intronization of gene open reading frames may mitigate endogenous silencing.

In summary, a comparative approach of epigenetic silencing has allowed us to dissect the biological rules underlying transgene silencing in these genomic contexts, which we exploit to develop a robust tool to achieve potent and selective gene reactivation of endogenously silenced transgenes in hiPSCs. This paves the way for the development of genetic anti-silencing modules that maintain transgene expression in circuits susceptible to silencing by endogenous cellular machinery. This is particularly relevant for dox-inducible systems that have widespread use, since it was shown that their inactivation was correlated with the level of DNA methylation at the minimal promoter ^21^, possibly due to reductions in transcription and H3K4me3 and subsequent recruitment of endogenous DNMTs, as mentioned above. Future work will explore the use of these anti-silencing modules in genomic integrations to ensure gene expression during stem cell differentiation and eventually, stem cell-derived cellular therapeutics. We anticipate that these methods will help advance cellular therapies that are more reliable and effective.

## Methods

### Cell Culture

All cells were maintained in a humidified incubator at 37°C with 5% CO2 under sterile conditions. CHO-K1 cells were cultured in tissue culture-treated 10 cm dishes (Corning) in Dulbecco’s Modified Eagle Medium (DMEM) (Corning) supplemented with 10% fetal bovine serum (FBS) (Corning), 1X non-essential amino acids (Gibco), and occasionally 100 U/ml penicillin (Sigma), and 100 µg/ml streptomycin (Sigma). Cells were passaged every 2-3 days. For regular passaging, the growth medium was aspirated and the cells were washed with 1X PBS without magnesium or calcium ions (Corning) before adding a sufficient amount of 0.25% trypsin-EDTA (Corning) and incubating at 37°C in a 5% CO2 incubator for 3-5 minutes to allow cell dissociation. The reaction was subsequently quenched by the addition of 1-2 volumes of complete DMEM. The cell suspension was transferred to 15 mL conical tube and centrifuged at 200 g for 3 minutes. After discarding the supernatant, the cell pellet was resuspended in fresh growth medium. The cell suspension was then diluted at a ratio of 1:10 - 1:20 and transferred to new tissue culture-treated plates with 12-14 mL of fresh complete medium. PGP1 human induced pluripotent stem cells (hiPSCs), generously provided by Prof. George Church at Harvard University, were cultured in mTesR Plus medium (STEMCELL Technologies) occasionally supplemented with 100 U/ml penicillin (Sigma), and 100 µg/ml streptomycin (Sigma). Medium was replaced every day or every other day depending on confluency. Cells were grown in 6-well plates pre-coated with hESC-qualified LDEV-free Matrigel (Corning). To coat the plates, Matrigel was diluted according to the dilution factor of the specific Lot Number in 25 mL of ice-cold DMEM/F12 before dispensing it in the culture plates at 1 mL/well for 6 well plates and 0.5 mL/well for 12-well plates. Plates were sealed with parafilm and stored at 4°C for up to 2 weeks until needed. Plates required for splitting or seeding were first incubated for at least 1 hour at room temperature or 30 minutes at 37°C to allow proper coating. Routine passaging of the hiPSCs was conducted every 3-5 days. Briefly, plates were first checked for differentiated regions, which were subsequently marked. Media was aspirated and differentiated regions were scraped off with a pipet tip. The remaining cells were washed with PBS and incubated with 1 mL Gentle Cell Dissociation Reagent (STEMCELL Technologies) for 4-6 minutes at room temperature. After aspirating the dissociation reagent, 1 mL pre-warmed mTeSR Plus medium was added, the colonies were detached using a sterile cell scraper and transferred to 15 mL conical tube and triturated 1-3 times with a 5 mL serological pipet. Finally, 1:10 - 1:40 of the population was reseeded into Matrigel-coated 6 well plates and placed in the incubator at 37°C with 5% CO2 overnight.

### DNA construction

The pX330-U6-Chimeric_BB-CBh-hSpCas9 plasmid used in this study was a gift from Feng Zhang (Addgene 42230). Target sgRNA sequences for the AAVS1 locus were designed using Bench-ling software. The DNA oligonucleotides AAVS1gRNAF and AAVS1-gRNAR were synthesized, annealed, and inserted into the pX330-U6-Chimeric_BB-CBh-hSpCas9 plasmid via Golden Gate assembly. The HDR donor plasmid was cloned by modifying the HDR template using restriction cloning. Sequences for chromatin regulators were cloned by hierarchical golden gate assembly. Briefly, epigenetic regulator sequences bearing kozak sequence and start / stop codons were ordered as gene fragments from Azenta and were flanked by BbsI cleavage sites and a 4bp buffer sequence. The gene fragment was inserted into a gene backbone by golden gate assembly using BbsI and T4 DNA ligase in T4 DNA ligase buffer (NEB) to obtain the level 0 part. The resulting plasmid comprises the epigenetic regulator ORF flanked by BsaI sites for modular cloning. The epigenetic regulator transcriptional unit was cloned by MoClo using the NEBridge GG BsaI Mastermix (NEB) and the resulting level 1 plasmid was purified using a Plasmid MIDI kit (Qiagen). Quantifications were performed using a Nanodrop 2000 after blanking with the supplied Qiagen elution buffer.

### Chromatin regulator design

We employed DNA binding domains of TetR, rTetR and PhlF for sequence-specific targeting ^122^. Effector domains used in this study include DNMT3A catalytic domain (UniProt Q9Y6K1 [aa602-912]), KRAB domain from ZNF10 (UniProt P21506 [aa11-75]), TET1 catalytic domain (UniProt Q8NFU7 [aa1419-2136]), dCas9-VPR (Addgene 63798), P300 core domain (UniProt Q09472 [aa1048-1664]), DOT1L catalytic domain (UniProt Q8TEK3 [aa1-416]), PRDM9 catalytic domain (UniProt Q9NQV7 [aa106-413]), EZH2 catalytic domain (UniProt Q15910 [aa503-727]), G9A catalytic domain (UniProt Q96KQ7 [aa829-1209]) and SUV91 catalytic domain (UniProt 43463 [aa76-412]). The KRAB domain was linked to rTetR or PhlF by GGGGS linker. All other effector domains were linked to DNA binding domain by flexible linker XTEN80 derived from E-XTEN [aa29-108] described by Schellenberger et al. ^123^. For nuclear localization, the nuclear localization sequence of the SV40 Large T-antigen (PPKKKRKV) was appended to the C terminal of each regulator. The full length fusion protein sequences are reported in Supplementary Information.

### Cell engineering

The CHO-K1 cell line used in this study was previously described in ^65^. To generate the hiPSC landing pad cell line (Fig. S3A), hiPSCs (PGP1) were transfected with 1000 ng of total DNA, maintaining a 1:1 ratio between the pX330-U6-Chimeric_BB-CBh-hSpCas9 plasmid with the inserted AAVS1 sgRNA and the donor plasmid. Following transfection, cells were cultured for 10 days in mTesR Plus medium, after which antibiotic selection was initiated by supplementing the medium with 200 µg/ml Hygromycin B (InvivoGen) for two weeks. To generate a single-cell suspension, the cells were incubated with Accutase (STEMCELL Technologies) for 8 minutes in a humidified incubator at 37°C with 5% CO2. The suspension was then centrifuged at 200 g for 3 minutes, and the resulting cell pellet was resuspended in mTesR medium supplemented with 10 µM Y-27632 and placed on ice. Fluorescence-activated cell sorting (FACS) was performed using a BD FACSAria to isolate EYFP-positive cells. The sorted cells were replated in Matrigel-coated plates in mTesR Plus medium containing 10 µM Y-27632 and 100 U/ml penicillin, 100 µg/ml streptomycin (Corning) for 24 hours, after which the Y-27632 supplement was withdrawn. A clonal cell line was then derived by seeding at clonal density in a Matrigel-coated 10 cm dish (50 cells/cm^2^ for 2,000-3,000 cells per dish) in mTesR Plus medium supplemented with 1X CloneR-2 and selecting individual colonies to expand.

For chromosomal integrations into the landing pad (Fig. S3A), hiPSCs bearing the landing pad were transfected with 1000 ng of DNA, maintaining a 1:1 ratio between the Bxb1 constitutive expression vector (CAG:Bxb1) and the reporter plasmid. At 24-48 hours post-transfection, antibiotic selection was initiated by feeding cells with fresh mTesR Plus medium supplemented with 1 µg/ml puromycin (InvivoGen) for a period of 7 days. At Day 10 post-trasnfection, a single-cell suspension was prepared by incubating the cells in Accutase (STEMCELL Technologies) for 8 minutes in a humidified incubator at 37°C with 5% CO2. The cell suspension was then trasnferred to a 15 mL contical tube and centrifuged at 200 g for 3 minutes, resuspended in mTesR Plus medium supplemented with 10 µM Y-27632 and 1 mg/mL BSA. Fluorescence-activated cell sorting (FACS) was conducted using a BD FACS Aria to isolate EBFP2+/EYFP-cells. Following sorting, the cells were cultured for 24-48 hours in mTesR Plus medium supplmented with 1X CloneR-2, 100 U/ml penicillin and 100 µg/ml streptomycin (Corning). Once viable colonies were visible, Cloner-2 supplement was withdrawn from the medium and the cells were expanded for banking. CHO-K1 cells were frozen as single cells in complete DMEM supplemented with 5% DMSO. PGP1 cells were frozen as aggregates in mFresR (STEMCELL Technologies). Freezing was performed in a Mr. Frosty to obtain a 1 °C/min cooling rate until −80°C, for a maximum time of 2 weeks short term storage at −80°C. Cells were transferred to a liquid nitrogen freezer for long-term storage.

### Transfections

Transfections were carried out in 12-well plates. For CHO-K1 cell transfections, they were treated with 0.25% trypsin-EDTA (Corning) and incubated for 3-5 minutes at 37°C with 5% CO2. Trypsin was neutralized by adding 1-2 volumes of pre-warmed complete growth medium at room temperature and the cell suspension was trasnfered to a 15 mL conical tube and at 200 g for 3 minutes. Then, the supernatant was aspirated and the cells were resuspended in sufficient complete growth medium before performing a viable cell count using Trypan Blue and a Countess (Invitrogen). Cells were seeded in 12-well tissue culture-treated plates (Corning) at a density of 125,000-150,000 cells per well and incubated for 24 hours in an incubator at 37°C with 5% CO2. The following day, the cells were washed once with 1X PBS and fed fresh growth medium free of antibiotics. Transfections were carried out using Lipofectamine LTX (Invitrogen) according to manufacturer’s instructions. Briefly, 800-1000 ng of DNA was combined in 50 µL of Opti-MEM (Gibco) with 1 µL of PLUS reagent in a 1.5 mL tube. Simlarly, 5 µL of Lipofectamine LTX was added to 50 µL of Opti-MEM in another 1.5 mL tube. The solutions were allowed to equilibrate for 5 minutes at room temperature, after which they were combined and incubated a further 5 minutes at room temperature to form the lipofection complexes. After incubation, 100 µL of formed transfection complexes were added dropwise to each well and evenly distributed by shaking. The media and trasnfection complexes were replaced after 24 hours in an incubator at 37°C with 5% CO2. For hiPSCs transfections, the cells were washed once with 0.5 mL 1X PBS and incubated in 0.5 mL Accutase (STEMCELL Technologies) for 8 minutes in a humidified incubator at 37°C with 5% CO2. Following incubation, 0.5 mL pre-warmed DMEM/F12 or PBS was added and the cell suspension was centrifuged at 200 g for 3 minutes, and the pellet was resuspended in an appropriate volume of fresh mTeSR Plus medium supplemented with 10 µM Y-27632 (STEMCELL Technologies) to obtain a single cell suspension. Viable cells were then counted using Trypan Blue and a Countess and seeded at a density 150,000-200,000 cells per well in Matrigel-coated 12-well plates in 1 ml mTeSR Plus supplemented with 10 µM Y-27632 (ROCKi). After 24 hours, Y-27632 was withdrawn from the medium. Transfections were carried out with Lipofectamine Stem (Invitrogen) according to manufacturer’s instructions. Briefly, 800-1000 ng of DNA was combined in 50 µL of Opti-MEM (Gibco) 1.5 mL tube. Simlarly, 4 µL of Lipofectamine Stem was added to 50 µL of Opti-MEM in another 1.5 mL tube. The solutions were allowed to equilibrate for 5 minutes at room temperature, after which they were combined and incubated a further 10 minutes at room temperature to form the lipofection complexes. After incubation, 100 µL of formed transfection complexes were added dropwise to each well and evenly distributed by shaking. The media and transfection complexes were replaced after 24 hours in an incubator at 37°C with 5% CO2.

### Cell sorting

CHO-K1 cells were detached by treatment with 0.25% trypsin-EDTA (Corning) at 37°C in a humidified incubator with 5% CO_2_ for 5 minutes. Trypsin was neutralized by the addition of 1 volume of complete growth media at room temperature. The cells were then centrifuged at 200 g for 3 minutes, after which the supernatant was discarded and the cell pellet was resuspended in complete growth media supplemented with 1 mg/ml bovine serum albumin (BSA) and the appropriate small molecules according to the experimental conditions (+/-2 µM dox). The cell suspension was filtered through a 35 µm filter cap into 5 mL round bottom polystyrene flow cytometry tubes. Fluorescence-activated cell sorting (FACS) was performed using a BD FACS Aria sorter 42-48 hours post-transfection, and 25,000-30,000 cells were collected in complete growth media supplemented with penicillin/streptomycin and the corresponding small molecules. Human iPSCs were detached with Accutase (STEMCELL Technologies) by incubating at 37°C in a humidified incubator with 5% CO_2_ for 7 minutes. The cells were then centrifuged at 200 g for 3 minutes, after which the supernatant was discarded, and the cell pellet was resuspended in mTeSR Plus supplemented with 10 *µ*M Y-27632, 1 mg/ml BSA and the appropriate small molecules according to the experimental conditions (2 µM dox or 30 µM DAPG). The cell suspension was filtered through a 35 µm filter cap into 5 mL polystyrene tubes for flow cytometry. Fluorescence-activated cell sorting (FACS) was performed using a BD FACS Aria sorter 42-48 hours post-transfection, and 25,000-30,000 cells were collected and cultured for 4 days on matrigel-coated plates in mTesR Plus supplemented 1X CloneR-2 (STEMCELL Technologies), penicillin/streptomycin and the corresponding small molecules based on the experiment. Fluorescence detection of EBFP2 was conducted using a 405 nm laser and a 450/50 filter with a voltage setting of 332 V, while EYFP was measured using a 488 nm laser and a 515/20 filter with a voltage setting of 200 V.

### Flow cytometry

CHO-K1 cells were detached by treating them with 0.25% trypsin-EDTA (Corning) and incubating at 37°C in a 5% CO2 incubator for 5 minutes. The reaction was quenched by adding 1 volume of complete warm growth medium. Cells were transferred to 1.5 mL tubes and centrifuged at 200 g for 3 minutes and the supernatant was aspirated. The resulting cell pellet was resuspended in an adequate volume of 1X PBS or fresh complete growth medium and passed through a 35 µm filter cap into 5 mL polypropylene tubes (Falcon) for flow cytometry analysis. Human iPSCs were detached with Accutase (STEMCELL Technologies) for 7 minutes in a humidified incubator at 37°C with 5% CO2. Following incubation, 1 volume of 1X PBS or DMEM/F12 was added and the suspension was transferred to 1.5 mL tubes before centrifugation at 200 g for 3 minutes. The supernatant was aspirated and the cell pellet was resuspended in 1X PBS or mTesR Plus medium supplemented with 10 µM Y-27632 and subsequently filtered through a 35 µm filter cap into 5 mL flow cytometry tubes (Falcon). Flow cytometry data was collected on a BD LSRFortessa, with specific laser and filter settings for each fluorophore: EBFP2 was detected using a 405 nm laser and a 450/50 filter at 259 V, EYFP with a 488 nm laser and a 530/30 filter at 190 V. Cytoflow (version 1.2) was utilized for data analysis, including calculating cell densities, means, standard deviations, and percentages from the flow cytometry measurements. The data were gated for single cells based on forward and side scatter, and only gated events were included in subsequent analysis. After data processing in Cytoflow, plots were generated using Cytoflow, GraphPad Prism, or Matplotlib. Bar plots were created using GraphPad Prism.

### CHO-K1 cell state switching

For transfections, 500 ng of rTetR-DNMT3A and 300 ng of EYFP were used for repression. Cells were sorted for high EYFP expression on day 2 after transfection and analyzed every 2 days by flow cytometry as described above. For reactivation, 500 ng of rTetR-TET1 and 300 ng of EYFP were used. Sorting and analysis was performed as before. Doxycycline was supplemented to the medium at a final concentration of 2 µM (1.03 µg/mL) for 8 days.

### CHO-K1 reactivation with chromatin regulators

For single transfection of a single regulator, 500 ng of the regulator and 300 ng of EYFP transfection marker were used. For co-transfections, 250 ng of either regulator and 300 ng of EYFP were used to maintain a total of 800 ng DNA. Doxycyline (2 µM) was supplemented to the medium for 4 days. Similarly, a 1:1 ratio was maintained between dCas9-VPR (250 ng) and the sgRNA (250 ng), with 300 ng EYFP. Cells were analyzed by flow cytometry as described above and computationally binned for EYFP expression.

### hiPSC silencing

For silencing, 500 ng of rTetR-DNMT3A or PhlF-KRAB along with 300 ng EYFP was used. In the case of rTetR-DNMT3A, 2 µM doxycycline was supplemented for 8 days. In the case of PhlF-KRAB, DAPG was added at a final concentration of 30 µM from days 8-12. Cells were sorted for medium high EYFP expression 2 days after transfection and analyzed by flow cytometry as described above.

### hiPSC reactivation

Reactivation of silenced hiPSCs was performed with 500 ng of rTetR-TET1 and 300 ng EYFP. Cells were sorted for medium high EYFP expression 2 days after trasnfection and analyzed by flow cytometry as described above.

### MeDIP-qPCR

MeDIP was conducted using the MagMeDIP-qPCR kit (Diagenode) following the manufacturer’s protocol with minor modifications. Transfections were performed as described above and cells were harvested following 10 days of culture. Genomic DNA was extracted from a confluent well of 12-well plate (approx. 3 million cells) using Monarch gDNA extraction kit (NEB T3010) according to kit instructions. Purified gDNA was eluted in 55 µL heated elution buffer. Quantification was performed on a Nanodrop, and DNA concentration was diluted to 100 ng/µL for those exceeding that value. DNA shearing was carried out in a Bioruptor Pico (Diagenode). For shearing, 50 µL of the DNA dilution was transferred to 0.2 mL Bioruptor tubes and sonicated for 30 cycles with a 30 s ON, 30 s OFF per cycle. Next, 47 µL of sheared DNA was denatured by combining with 10 µL ultrapure water and 33 µL 1X MagMaster Mix for 3 minutes at 95 °C according to manufacturer protocol and then cooled quickly on ice for 10 minutes. For antibody-bead binding, 75 µL of denatured sheared DNA was combined with 20 µL of washed MagBeads and 5 µL of antibody mix according to manufacturer protocol. The anti-5mC antibody Clone 33D3 (Diagenode - C15200081) was used. The DNA-bead-antibody solution was rotated overnight at 4 °C. The following day, beads were washed 4 times with the supplied wash buffers as per instructions (3x 100 µL buffer A, 1x 100 µL buffer B) and the enriched DNA was eluted in 100 µL of DNA isolation buffer supplemented with Proteinase K by incubating for 15 minutes at 55 °C then 15 minutes at 100 °C. The supernatant was recovered from the beads using a magnetic rack and used as input for qPCR reactions. All qPCR reactions were performed in technical duplicates using Power SYBR Green PCR Master Mix (Applied Biosystems) and primers on a LightCycler 96 system (Roche). Reactions consisted of 50 µL final volume, with 10 µL input DNA and 0.4 µM forward and reverse primers. Melt curve analysis was performed to verify a single amplification product, and the LightCycler 96 Application Software (Roche) was used to obtain Ct values. The promoter region of the reporter gene was the target location to be amplified. IgF2 in CHO-K1 and TSH2B in human cells served as reference endogenous genes. The fold changes reported were calculated using the standard *δδ*Ct method. Primer sequences are reported in Supplementary Information.

### CUT&RUN-qPCR

CUT&RUN-qPCR was performed according to ^124^ with minor modifications. Transfections were performed as described above and cells were harvested following 10 days of culture. Cells were harvested (CHO-K1 cells with 0.25% Trypsin-EDTA for 3-5 minutes, PGP1 cells with Acctuase for 8-10 minutes), washed twice with 0.5 mL 1X PBS by centrifuging at 600 g for 3 minutes and discarding supernatant and finally resuspending in 1 mL PBS. Next, a viable cell count with Trypan blue exclusion was performed and 1 million cells per condition were transferred to fresh 1.5 mL tubes and pelleted by cetrifugation at 600 g for 3 minutes. The resulting pellet was washed twice with 1 mL wash buffer (20 mM HEPES pH 7.5, 150 mM NaCl, 0.5 mM spermidine, 1X EDTA-free protease inhibitor) and resuspended in 990 mL of wash buffer. Meanwhile, 100 µL of Concanavalin A Magnetic beads (Bangs Labs - BP531) were washed three times in 1 mL binding buffer (20 mM HEPES pH 7.5, 20 mM KCl, 1 mM CaCl_2_, 1 mM MnCl_2_), and finally resuspended in 100 µL of binding buffer (enough for 9 samples, scale accordingly, 10 µL beads per sample + 10% excess). For cell-bead binding, 10 µL of the washed beads were added to the 990 mL of washed cells and rotated at room temperature for 15 minutes. Next, the supernatant was discarded and the beads-cells were resuspended in 200 µL of antibody buffer (wash buffer + 0.05% digitonin + 2 mM EDTA) and 3 µL of the antibody was added. We used a polyclonal antibody specific to histone 3 lysine 9 trimethylation (H3K9me3) (Active Motif - 39161). The cell-bead-antibody slurry was then transferred to 0.2 mL PCR tubes and rotated overnight at 4 °C. The following day, the beads were washed twice with 200 µL of digitonin buffer (wash buffer + 0.05% digitonin), then resuspended in 200 µL digitonin buffer and 2.5 µL of pAG-MNase (Active Motif - 53181) was added. The tubes were rotated for 30 mintutes at room temperature. Next, the beads were washed twice with 200 µL of digitonin buffer before being resuspended in 200 µL digitonin buffer and cooled on an aluminum rack on ice for 10 minutes. To initiate chromatin digestion, 2 µL of 0.1 M CaCl_2_ was added to the cooled bead slurry, combined well and rotated for 4 hours at 4 °C in a cold room. Chromatin digestion was terminated by the addition of 67 µL of stop buffer (5 mM NaCl, 20 mM EDTA, 4 mM EGTA, 50 µg/mL RNase A, 50 µg/mL Glycogen). The tubes were then incubated in a thermocycler at 37 °C (lid 65 °C) for 20 minutes, then spun on a tabletop micro-centrifuge, placed on magnetic rack and the supernatant (aprox. 267 µL - sample) was transferred to fresh 2.0 mL tubes for purification with Monarch Spin PCR&DNA cleanup kit (NEB - T1130). Briefly, 5 volumes (1335 µL) of binding buffer were added to the recovered supernatant, passed through the supplied spin tubes by centrifugation, washed twice with 500 µL of 80% ethanol and eluted in 20 µL elution buffer. The eluate was used as the DNA input for qPCR reactions. All qPCR reactions were performed in technical duplicates using Power SYBR Green PCR Master Mix (Applied Biosystems) and primers on a LightCycler 96 system (Roche). Reactions consisted of 50 µL final volume, with 3 µL input DNA and 0.4 µM forward and reverse primers. Melt curve analysis was performed to verify a single amplification product, and the LightCycler 96 Application Software (Roche) was used to obtain Ct values. The promoter region of the reporter gene was the target location to be amplified. IgF2 in CHO-K1 and TSH2B in human cells served as reference endogenous genes. The fold changes reported were calculated using the standard *δδ*Ct method. Primer sequences are reported in Supplementary Information.

### Immunostaining for flow cytometry

Cells were prepared for staining by first incubating them in Accutase (STEM CELL Technologies) for 7 minutes. After incubation, the cells were resuspended in 4 volumes of mTesR medium and centrifuged at 200 g for 3 minutes. The pellet was resuspended in mTeSR containing 10 µM ROCK inhibitor (Y-27632), followed by another centrifugation at 200 g. The cells were then resuspended in phosphate-buffered saline (PBS) and washed twice with PBS, each wash involving centrifugation at 200 g for 3 minutes. Following the final wash, the cells were resuspended in 100 µL of staining solution containing anti-TRA-1-60-AF647 antibody diluted 1:50 in PBS. The staining was performed in a 96-well plate, with tissue culture-treated wells used for human induced pluripotent stem cells (hiPSCs) and non-tissue culture-treated wells for CHO-K1 cells. The cells were incubated overnight for 18 hours at 4°C, protected from light by covering the plates with aluminum foil. The following day, the cells were washed twice with PBS to remove excess antibody, and resuspended in 250 µL of PBS for flow cytometry analysis.

### Microscopy

Imaging was performed using a Zeiss Axio Observer Z1 inverted widefield fluorescence microscope equipped with a AxioCam MRm camera and X-Cite Series 120 light source in a humidified chamber at 37 °C and 5%CO_2_. Images were acquired on ZenPro software with objective A Pln 20x/0.8 DICII in bright field and the EBFP2 fluorescence channel. In total, 9 tiles (3×3 images) were acquired and stitched automatically by the software. Contrast adjustments and merged images were generated with MATLAB.

### Bulk RNAseq

Cell seeding, transfection and sorting was performed as described above. In particular, 500 ng of TetR-TET1 and 300 ng EYFP were used for transfection. Sorting was performed 2 days later, and populations in the active state, endogenously -silenced state as well as those with high trasnfection marker were sorted and maintained in culture for 8 days. On day 10, cells were treated with Accutase for 8 mins at 37°C, washed twice with 1X PBS and viable cells were counted. Total RNA was extracted from 3 million cells with the Monarch Spin RNA Isolation Kit (NEB - T2110S) and eluted in 100 µL Ultrapure water before quantification on a Nanodrop. Samples were submitted to the MIT BioMicro Center for sequencing. Briefly, mRNA was enriched with poly-A bead capture, adapters were ligated to form the library and sequencing was performed on a SingularG4 sequencer. Genomic sequences, transcriptomic sequences, and gene annotation files were obtained from GENCODE (release 49). The EBFP2 sequence was then appended to each of the obtained files. Genome indexing and paired-end sequence alignment was performed with bowtie (version 1.2.3) ^125^. Aligned sequences were counted using the featureCounts function from the Rsubread R package (version 2.12.3) ^126^. Transcriptome indexing and transcript quantification was performed with salmon (version 1.10.0) ^127^. Differential expression analyses were performed using DESeq2 (version 1.38.3) ^128^. Only genes with at least 10 counts in at least 3 samples were included in the differential expression analysis. Raw data was deposited on Harvard Dataverse and can be accessed at https://doi.org/10.7910/DVN/FKA79G.

### Targeted TAPS+ Sequencing

Genomic DNA was extracted using the Monarch genomic DNA extraction kit (NEB) and eluted in 50 uL TE buffer. DNA conversion and sequencing was performed by the BioMicro center at MIT. Briefly, TAPS+ conversion (Watchmaker Kit) was performed according to manufacturer’s instructions with minor modifications. Genomic DNA (100 ng) was oxidized and reduced following kit protocol, and the DNA subsequently amplified with 2 primer pairs targeting the operator and promoter sequences, where the first 6 cycles were performed with DHU polymerase (Watchmaker) for conversion of modified C bases to T, after which the DNA was further amplified with Q5 polymerase (NEB). The 5’ end of the primers contained a landing sequence (forward primers have “ACACGACGCTCTTCCGATCT” and the reverse primers have “CGGCATTCCT-GCTGAACCGCTCTTCCGATCT”) for the second PCR with the Q5 polymerase. The resulting amplicons were barcoded, pooled and sequenced on an Oxford Nanopore Promethion P2. Raw data was deposited on Harvard Dataverse and can be accessed at https://doi.org/10.7910/DVN/UVODUS.

### Lentiviral transductions

Lentivirus was produced with Lenti-X Packaging Single-Shot VSV-G preparation by transfecting 293T cells grown in DMEM supplemented with 10% FBS with Packaging plasmids and donor plasmids. The supernatant was collected 72 hours following transfection (aprox. 8 mL) and concentrated with Lenti-X concentrator (to aprox. 1.5 mL). PGP1 cells were transduced in mTeSR Plus supplemented with CloneR-2. CHO-K1 cells were transduced in DMEM supplemented with 10% FBS and 1% NEAA. Reverse transductions were carried out with 100,000 cells, 100 µL of concentrated virus and 8 µg/mL PolyBrene in 1 mL of medium in a 6-well plate for 24 hours. Cells were sorted by FACS 72 hours post-transduction for EBFP2 expression, generating polyclonal populations from 2000 cells for PGP1 and 5000 cells for CHO-K1. Chromatin editing of lentivirus-generated cell lines was performed similar to the experiments detailed above.

### Computational analysis

All simulations conducted in this paper were done using Gillespie’s Stochastic Simulation Algorithm (SSA) ^129^.

## Acknowledgements

We thank the Koch Institute’s Flow Cytometry Core at MIT for the support with cell sorting. We thank the MIT BioMicro Center for their assistance with RNAseq data acquisition as well as the TAPS+ methyl CpG sequencing and analysis. Furthermore, we would like to thank our funding: NSF-MODULUS (Award number 2027949) and NIH-NIBIB (Award number R56EB036090).

## Supplementary Information (SI)

### 1 Supplementary Text

#### Chromatin modification model with DNA methylation and H3K9me3

In this section, we describe the chemical reaction and ODE model corresponding to the reaction diagram of Fig. 3, which describes the interactions between DNA methylation and H3K9me3 consistent with our data and with the literature. To this end, we represent by U nucleosomes devoid of both DNA methylation and H3K9me3, we let x represent nucleosomes modified with H3K9me3 only, y nucleosomes modified with DNA methylation only, and z nucleosomes modified with both H3K9me3 and DNA methylation. We further let K denote the KRAB input, which recruits enzymes that write H3K9me3, let D represent DNMT3A, and let T represent the TET1 enzyme. With these definitions, the chemical reactions corresponding to the reaction diagram are as follows.

The establishment of H3K9me3 occurs either *de novo* through K or through the autocatalysis by nucleosomes that have H3K9me3^93–95^:

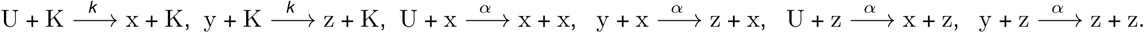

Additionally, in our experiments H3K9me3 can be brought about by DNA methylation (Fig. 2F), which is consistent with the literature reporting that DNA methylation recruits writers of H3K9me3^90^. To account for this in our model, we add reactions where nucleosomes with DNA methylation (y or z) recruit H3K9me3 on histones devoid of it (U and y):

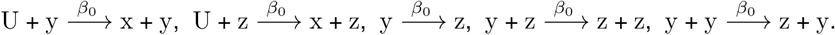

The establishment of DNA methylation also occurs through two different pathways, that is, *de novo* establishment through D on nucleosomes devoid of DNA methylation (U and x):

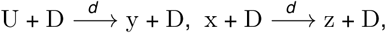

or via the recruitment of DNMT3s by H3K9me3 (x or z) to histones lacking DNA methylation (U or x), consistent with our data (Fig. 2F) and the literature ^91,92^:

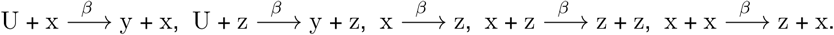

In cell lines lacking either DNMT3s or reader proteins that bind to H3K9me3 and recruit DNMT3s, we have that *β* = 0.

Finally, we have the decay of H3K9me3, with rate constant *δ*:

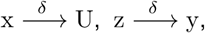

and the decay of DNA methylation, which is non-negligible only in the presence of the TET1 enzyme T, consistent with our data (Fig. 2F) and the literature ^96,97^:

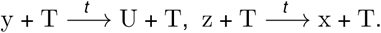

We let *x* denote the fraction of nucleosomes with H3K9me3, *y* the fraction of nucleosomes with DNA methylation, *z* the fraction of nucleosomes with both modifications, and *u* the fraction of nucleosomes with no modifications. By lumping the *t*, *k*, and *d* rate constants into the level of inputs *T* for TET1, *K* for KRAB, and *D* for DNMT3A, we obtain the following system of ordinary differential equations (ODEs) describing the rate of change of these fractions:

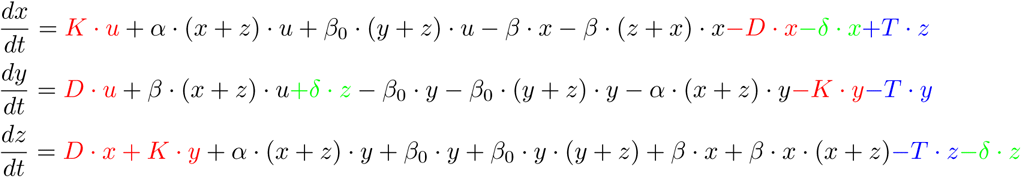

in which *u* = 1 − *x* − *y* − *z*. The total amount of H3K9me3 *x̄* = *x* + *z* obeys the differential equation:

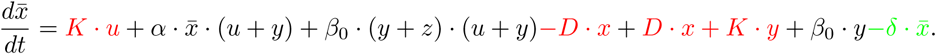

When *T* is very large and no inputs *D* or *K* are present, *y* ≈ 0 and *z* ≈ 0, which leads to the simplified *x̄* dynamics as:

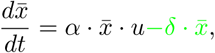

in which *x̄* ≈ *x* and *u* ≈ 1 − *x*, which is the same ODE as in Fig. 3B. The simulations in Fig. 3DE are obtained with the following parameter values: *α* = {0.1, 10}, *β* = 0.1, *β*_0_ = 0.1, *δ* = 1, *T* = 10, *D* = 1, and *K* = 1. Note that we have let *β* and *β*_0_ to be smaller than *α* in the simulations. In fact, if these two parameters are too large, the full system’s positive feedback around H3K9me3 and DNA methylation will cause a small residual DNA methylation to push up H3K9me3 levels, which in turn will re-establish DNA methylation, and, similarly, small residual H3K9me3 can push up DNA methylation levels, which re-establish H3K9me3 (Fig. S9AB).

### 2 Supplementary Tables

**Table 1:**
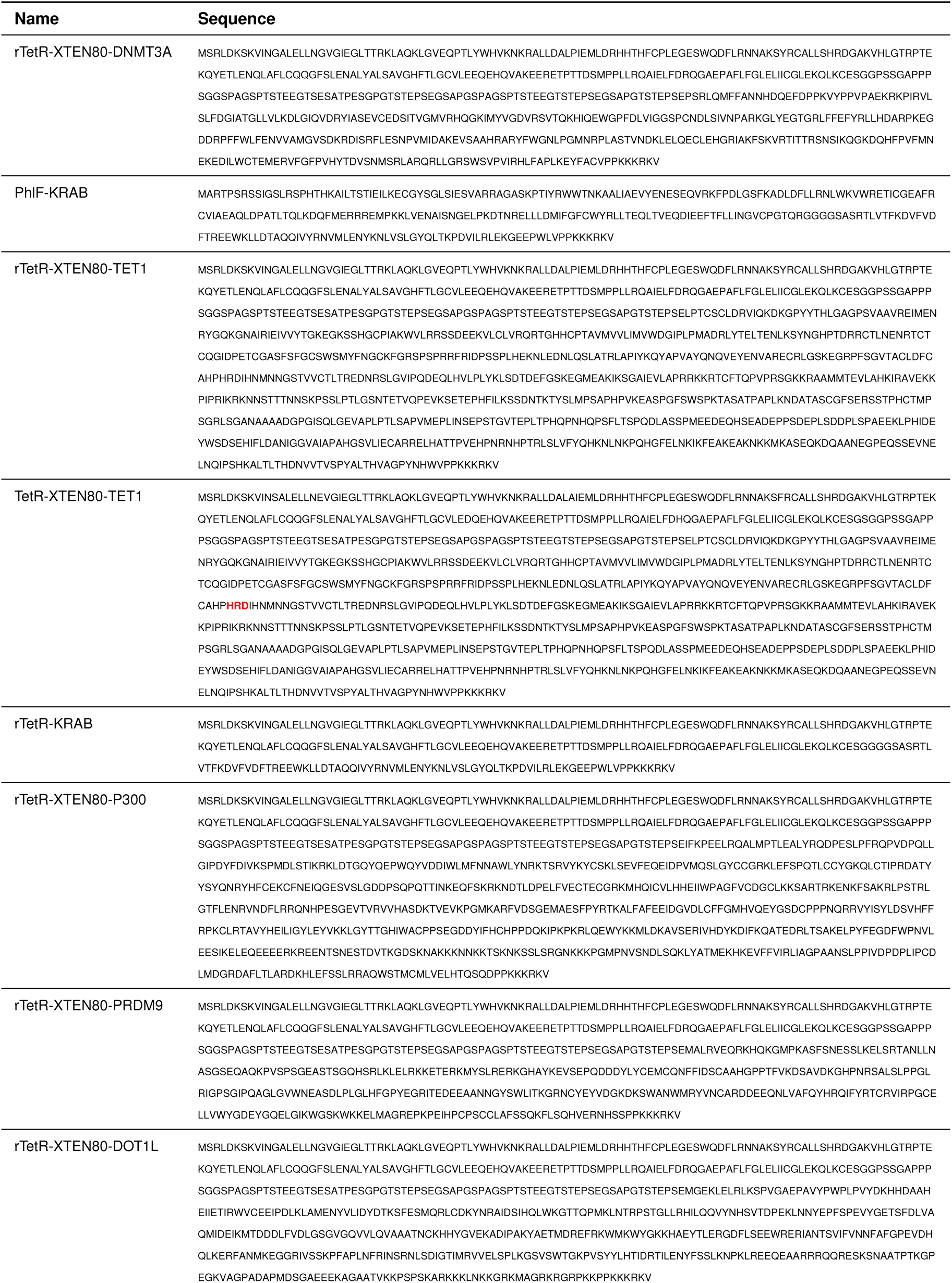
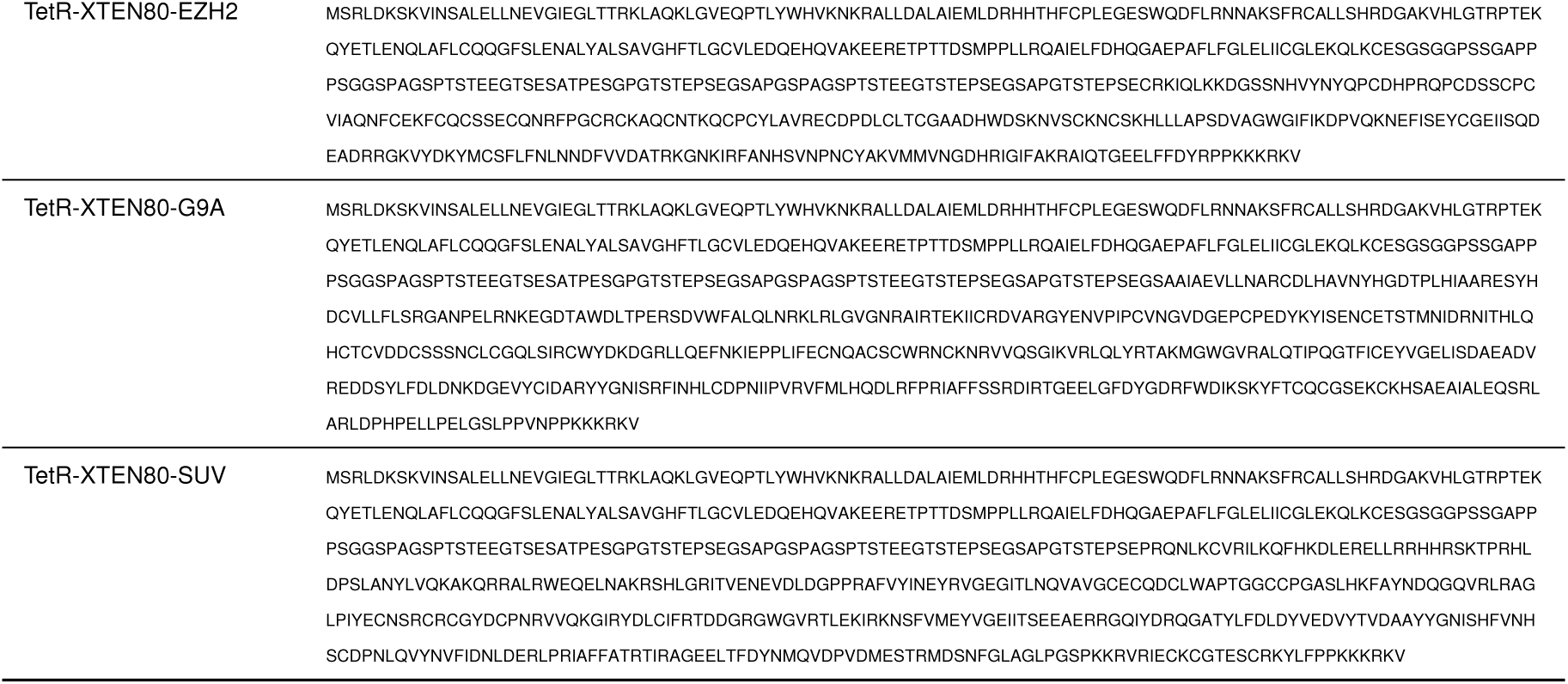
Chromatin regulator sequences. The bold red residues (HRD) in TetR-TET1 are mutated to YRA in TetR-deadTET1.

**Table 2:**
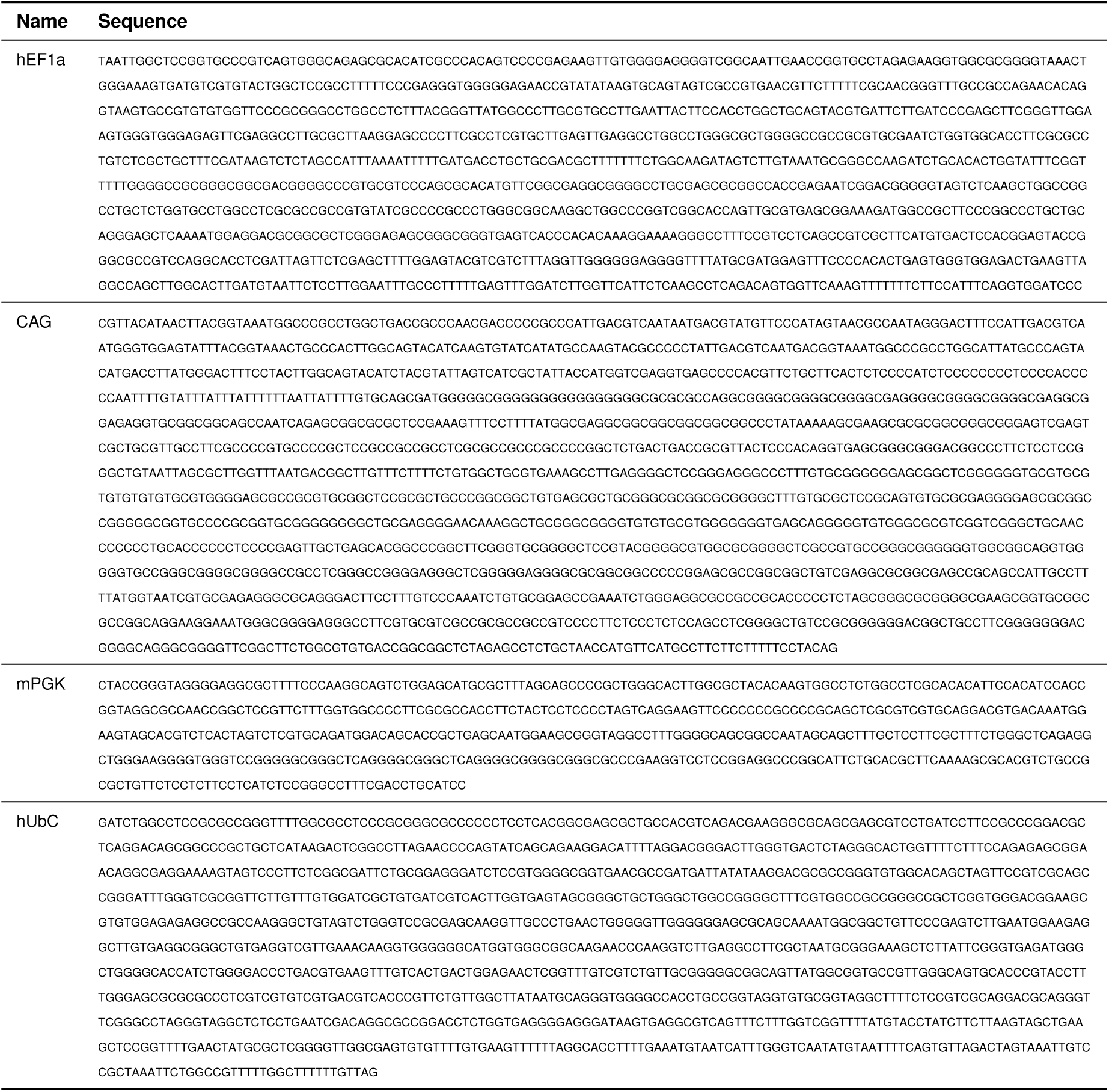
DNA sequences.

**Table 3:**
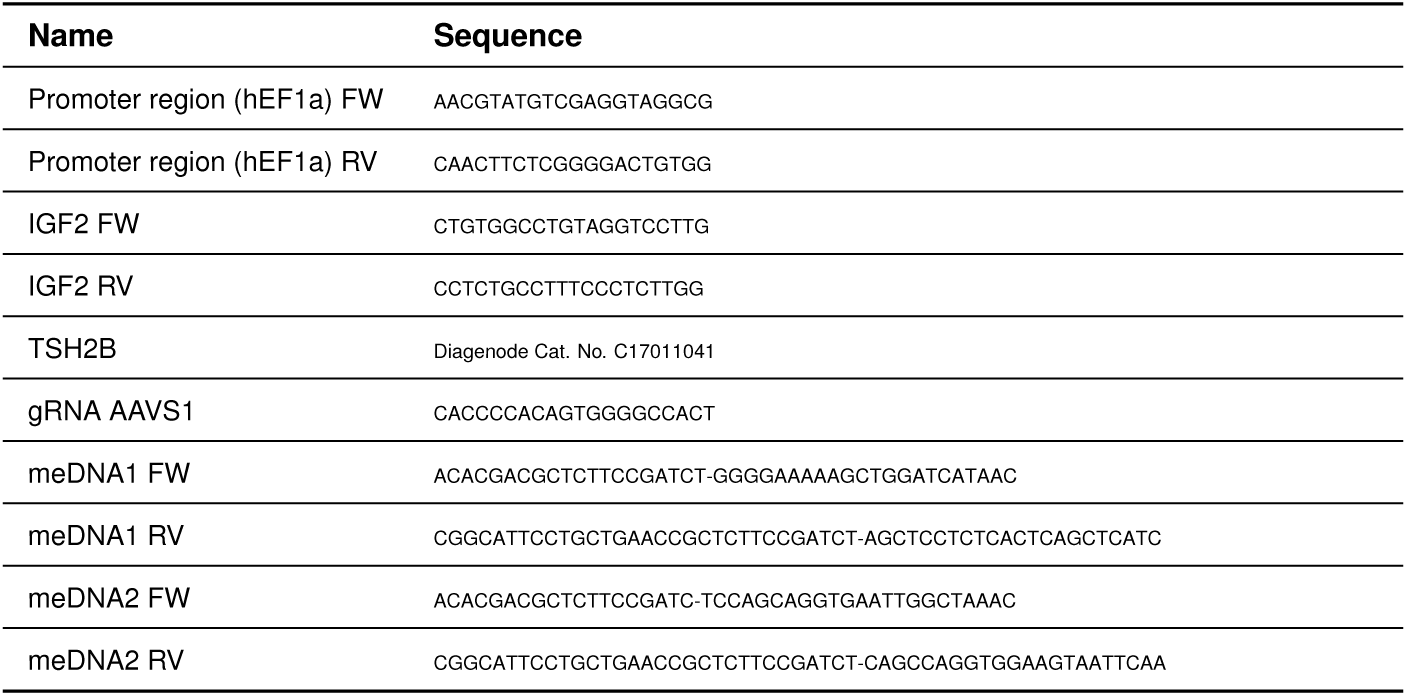
Primer and gRNA sequences (5’-3’) Note that 5’ end of meDNA primers contain a landing sequence (before the hyphen) for a second PCR amplification with Q5 polymerase after DHU Polymerase conversion (see Methods).

### 3 Supplementary Methods

Cell culture, seeding, transfection, sorting and flow analysis were performed as described in Main Methods. The following methods describe the amounts of DNA transfected for experiments appearing in the Supplementary Figures.

**Comparing DNMT3A and KRAB silencing in CHO-K1 cells.** 500 ng of rTetR-DNMT3A or PhlF-KRAB was transfected with 300 ng EYFP marker with Lipofectamine LTX to 150,000 cells in a 12 well plate seeded the day before. Sorting was performed on day 2, and flow cytometry measurements were acquired after sorting and every other day thereafter until day 8.

**Non-targeted chromatin regulators.** 500 ng of DNMT3A or KRAB without fusion to DNA binding domain was transfected along with 300 ng EYFP as described in Methods. On day 5 after transfection, cells were analyzed by flow cytometry. The data were computationally binned for EYFP marker expression and plotted accordingly.

**TET1 or rTetR for reactivation.** 500 ng of TET1 without a DNA binding domain or 500 ng of rTetR without a catalytic domain were transfected along with 300 ng EYFP marker. On day 4 after transfection, cells were analyzed by flow cytometry. The data were computationally binned for EYFP marker expression and plotted accordingly. For rTetR transfections, 2 µM doxycycline was used throughout the experiment.

**Silencing and reactivation of various promoters in CHO-K1 cells.** For silencing, 500 ng rTetR-DNMT3A and 300 ng EYFP were transfected. Doxycycline was administered for 8 days. For reactivation, 500 ng rTetR-TET1 and 300 ng EYFP were used. Dox was administered for 6 days. The final dox concentration was 2 µM.

**Silencing of CAG promoter reporter in hiPSCs.** For single transfection, either 500 ng of rTetr-DNMT3A or 500 ng rTetR-KRAB were used along with 300 ng EYFP marker. For co-transfection of both chromatin regulators, 250 ng of each was transfected with 300 ng of EYFP. The cells were sorted 2 days later and monitored for 19 days. Dox (2 µM) was supplemented for 8 days after transfection.

**Reactivation of silenced CAG reporter in hiPSCs.** To obtain the repressed starting population, cells in the silenced state from the previous rTetR-DNMT3A and rTetR-KRAB silencing were sorted for EBFP2- and expanded. For TET1 transfections, 500 ng of TetR-TET1 and 300 ng EYFP marker were used. For VPR transfections, 250 ng of each dCas9-VPR and gRNA targeting TetO with 300 ng EYFP were used. Cells were sorted 2 days after transfections and cultured for 15 days. Flow cytometry analysis was performed after sorting on day 2, then day 7 and 15.

### 4 Supplementary Figures

**Fig. S1.**
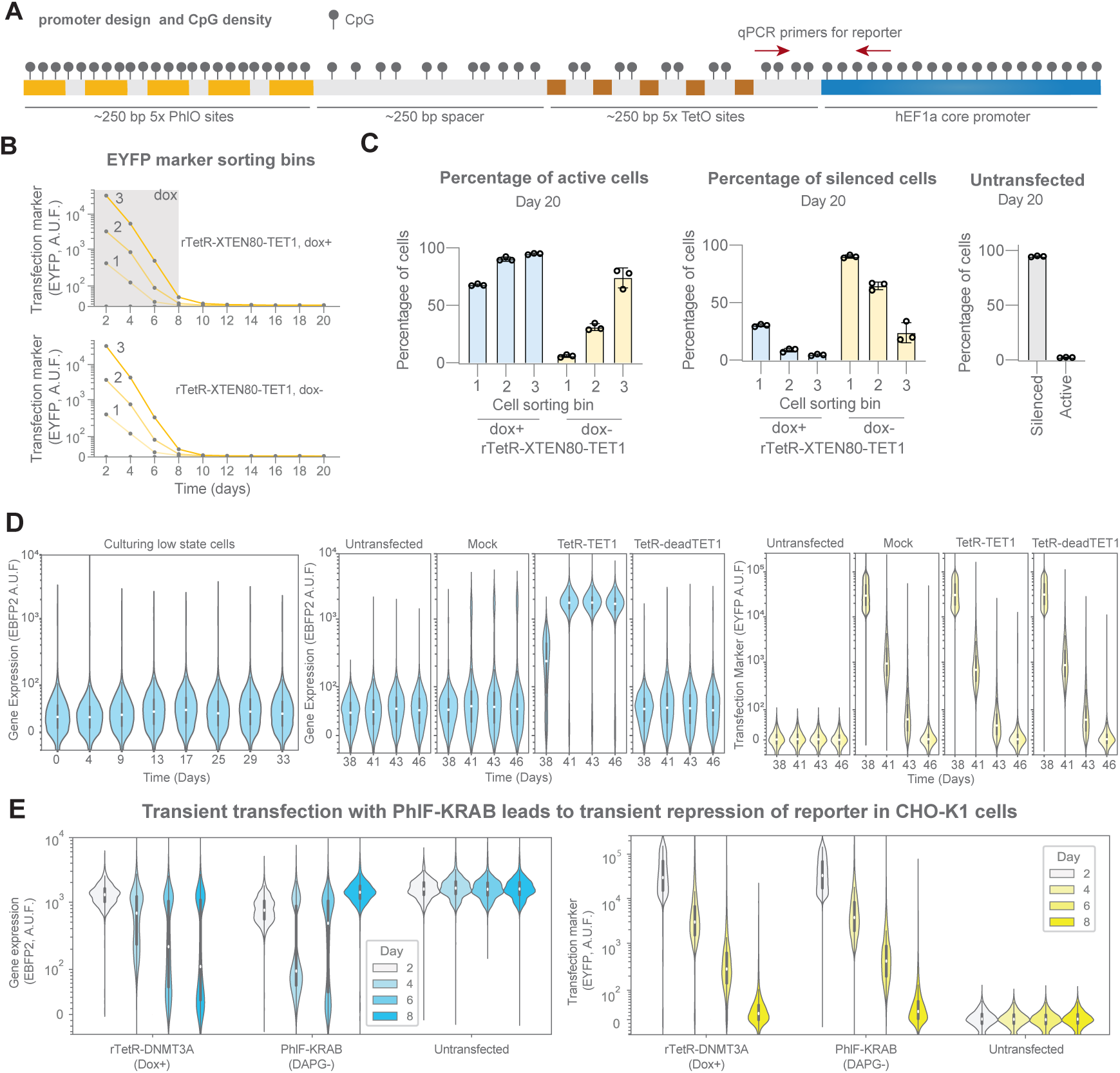
Chromatin regulation in CHO-K1 cells. **A.** Design of operator-promoter pair used to drive reporter gene expression and distribution of CpG sites. Not to scale. The hEF1a promoter intron is not shown. This panel is related to Fig. 1A**. B.** Scatter plot of the flow cytometry median EYFP values for the different cell sorting bins (Bin 1 = 10^2^ *−* 10^3^, Bin 2 = 10^3^ *−* 10^4^, Bin 3 = 10^4^ *−* 10^5^ AUF) over time for reactivation of CHO-K1 cells by transient rTetR-TET1 transfection. This panel is related to Fig. S1C. **C.** Bar plots showing the percentage percentages of active and silenced cells at day 20 after transfection with rTetR-TET1 (dox+/-) for the different sorting bins shown in S1B. **D.** Violin plots showing reporter gene expression (EBFP2) after extended culture time in the silenced state (left), reporter gene expression after trasnient transfection of Mock, TetR-TET1 or TetR-deadTET1 (center) and their correlated transfection marker expression levels (EYFP, right). Mock transfection is a emplty plasmid backbone, deadTET1 is a catalytically inactive TET1 domain. **E.** Violin plots showing reporter gene expression (EBFP2) and transfection marker expression (EYFP) after transient transfection with chromatin regulators rTetR-DNMT3A or PhlF-KRAB in CHO-K1 cells over 8 days. Data are plotted from three independent replicates. For all violin plots, the white dot represents the median, the thick box represents the interquartile range (IQR) and the thin gray line represents 1.5x the IQR.

**Fig. S2.**
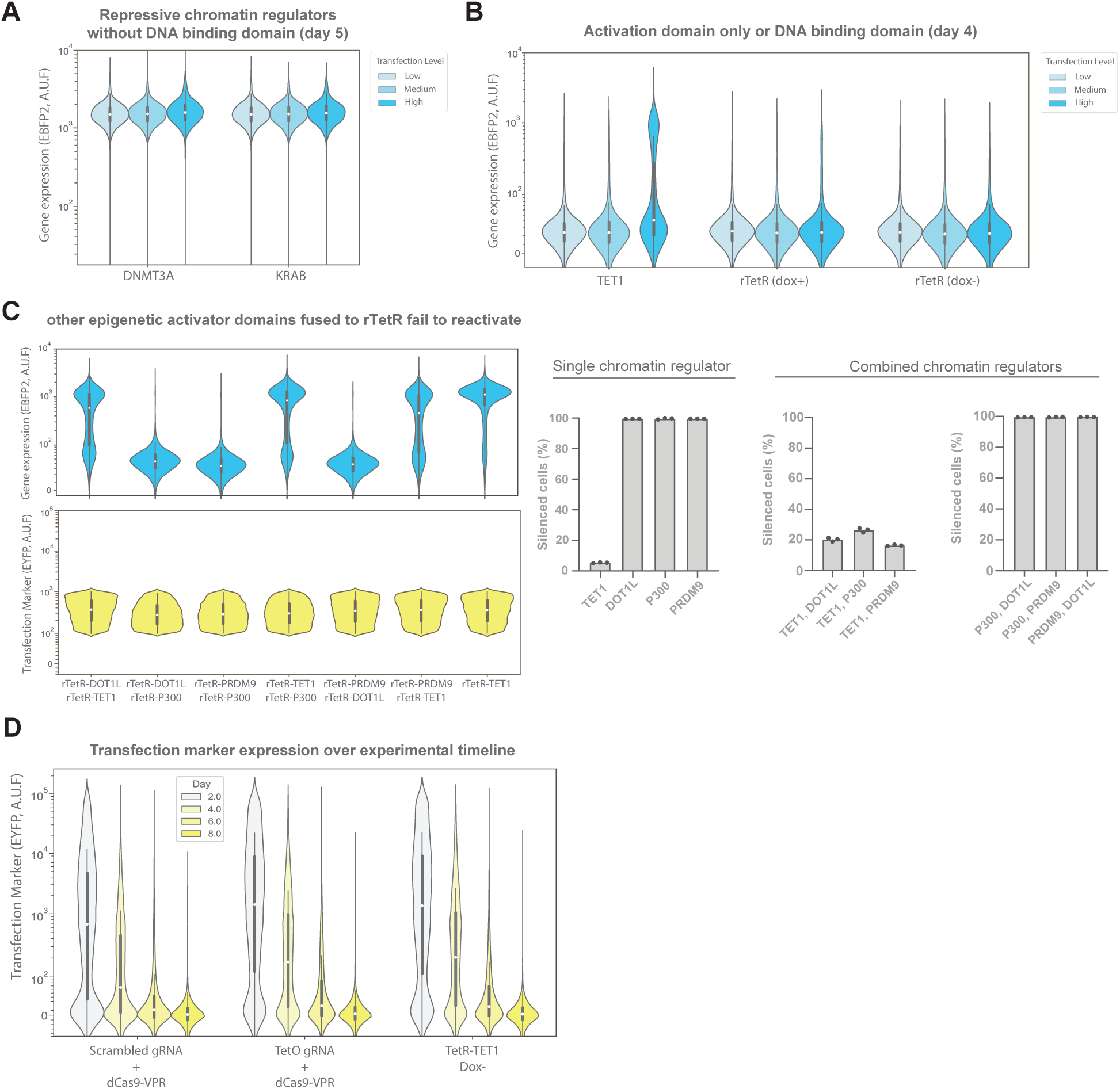
Non-targeting chromatin regulators and transfection marker levels. **A.** Violin plots showing transfection of repressive chromatin regulators without DNA binding domain. **B.** Violin plots showing transfection of activation domain (TET1) without DNA binding domain, or DNA binding domain only (rTetR) in presence or absence of doxycycline (dox). **C.** Violin plots showing reporter gene expression (EBFP2) and transfection marker expression (EYFP) for transient co-transfection of various activating chromatin regulators. Bar plots (right) show the fraction of silenced cells estimated from the violin plots for the different conditions. This panel is related to Fig. 1E. **D.** Violin plots of transfection marker expression for dCas9-VPR and TetR-TET1 modulation comparison. This panel is related to Fig. 1F. For all violin plots, data are plotted from three independent replicates. The white dot represents the median, the thick box represents the interquartile range (IQR) and the thin gray line represents 1.5x the IQR.

**Fig. S3.**
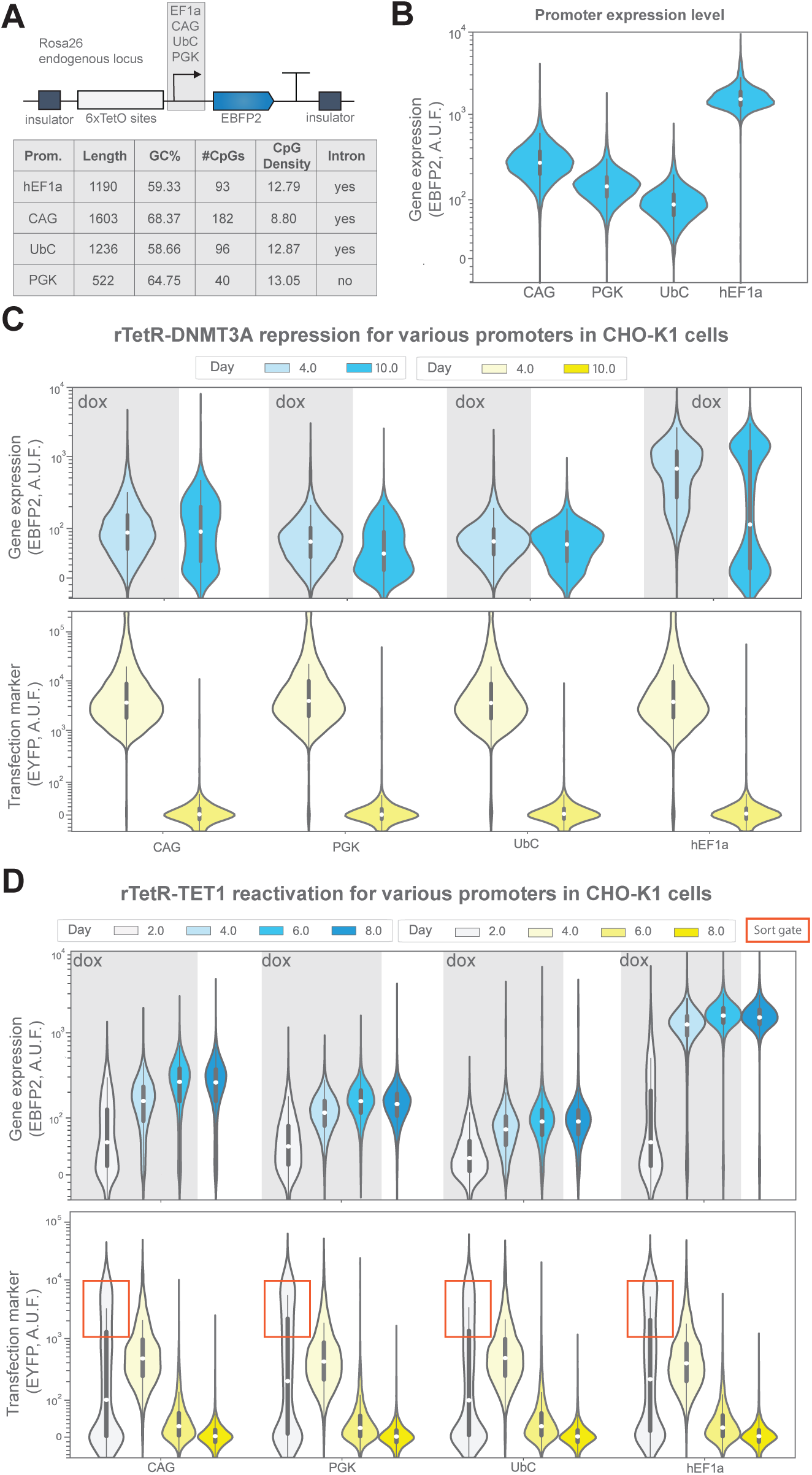
Comparison of mammalian constitutive promoters for silencing and reactivation dynamics in CHO-K1 cells. **A.** Schematic showing reporter construct with the various constitutive promoters employed (CAG, mPGK, hUbC and hEF1a) and their relevant properties. **B.** Violin plots comparing promoter output of EBFP2 fluorescence following site-specific integration. **C.** Violin plots comparing silencing dynamics of the various promoters following transient rTetR-DNMT3A transfection. **D.** Violin plots comparing reactivation dynamics of the various promoters following transient rTetR-TET1 transfection. For all violin plots, data are plotted from three independent replicates. The white dot represents the median, the thick box represents the interquartile range (IQR) and the thin gray line represents 1.5x the IQR.

**Fig. S4.**
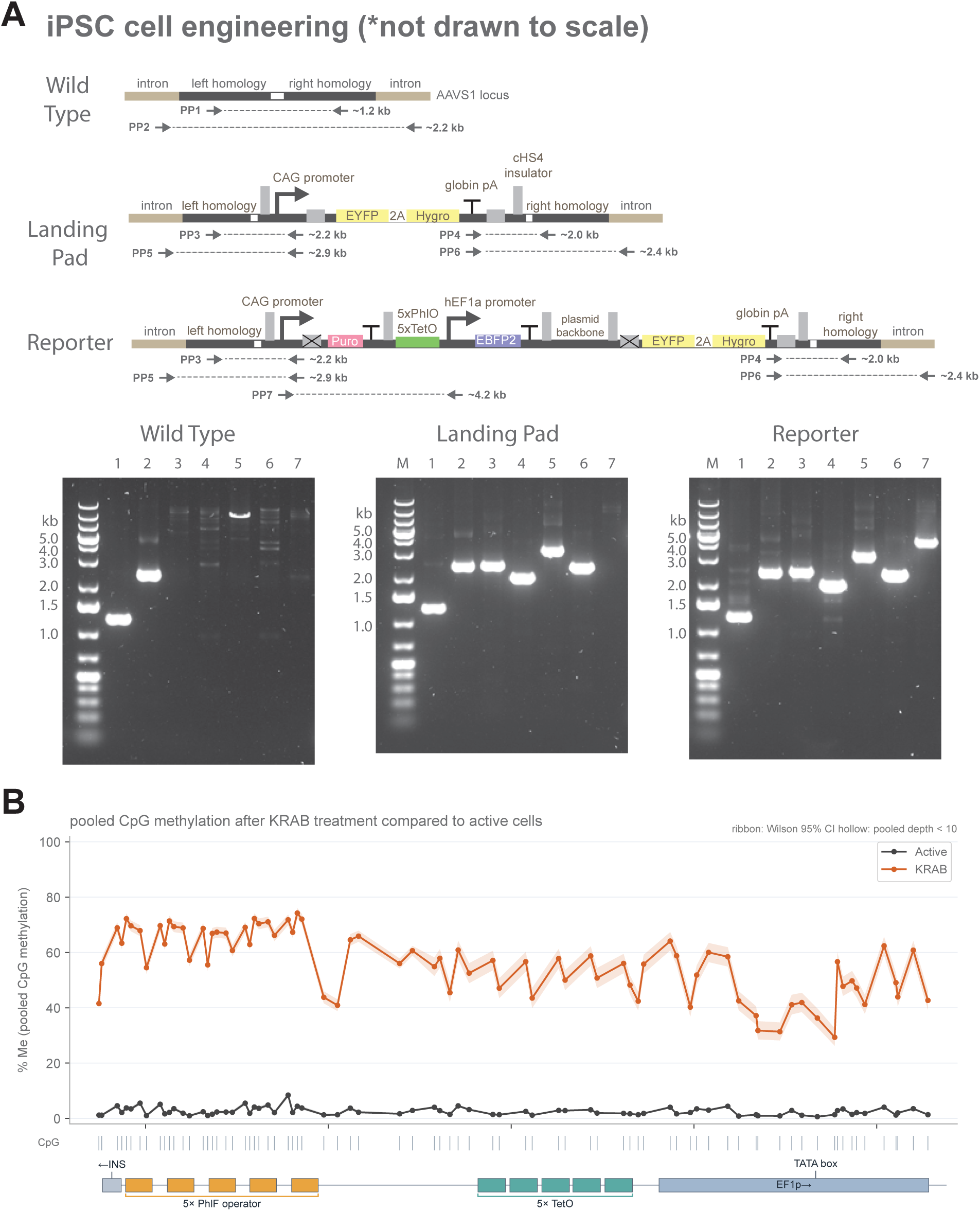
iPS cell engineering and methylation analysis after KRAB treatment. **A.** Schematic showing design generation of landing pad cell line followed by the reporter line. Primer Pairs (PP shown as arrows) were used to verify location site and integrated DNA. Marker (M) is the Generuler 1 Kb plus (invitrogen). This panel is related to Fig. 2A. See Methods for details regarding integration. **B.** Line plot showing CpG methylation levels of the transgene operator-promoter pair for cells in the active state and after targeted KRAB treamtent. Data are average from three independent replicates and shaded error represents 95% confidence interval.

**Fig. S5.**
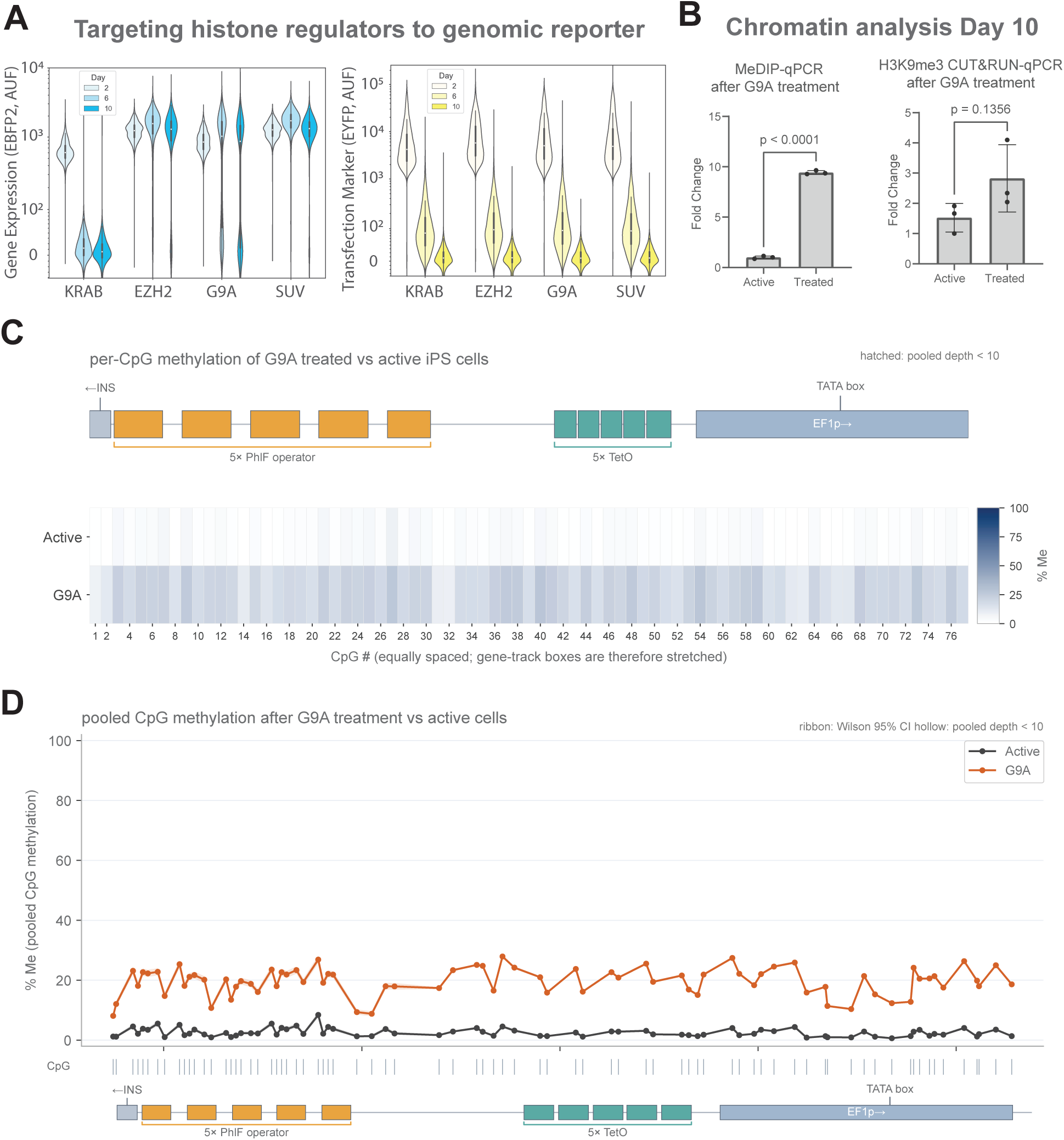
Targeted H3K9me3 deposition leads to recruitment of DNA methylation in hiPSCs. **A.** Violin plots showing gene expression leveles (EBFP2) and transfection marker levels (EYFP) after targeted silencing of the reporter with histone methylases (fused to TetR). **B.** Quantitative PCR analysis (qPCR) of chromatin after methylated DNA immunoprecipitation (MeDIP) or CUT&RUN targeting H3K9me3 at Day 10 after TetR-G9A transfection. Bars represent mean values of three independent replicates. Statistics were performed with an ordinary student’s T-test for pairwise comparison. **C.** Heatmap of targeted TAPS+ sequencing of the reporter transgene promoter. Methylation level of each CpG residue within the operator and promoter regions are shown for active and G9A-silenced cells. Data reported are average from three independent replicates. **D.** Line plot showing CpG methylation levels of the transgene operator-promoter pair for cells in the active state and after targeted G9A treamtent. Data are average from three independent replicates and shaded error represents 95% confidence interval.

**Fig. S6.**
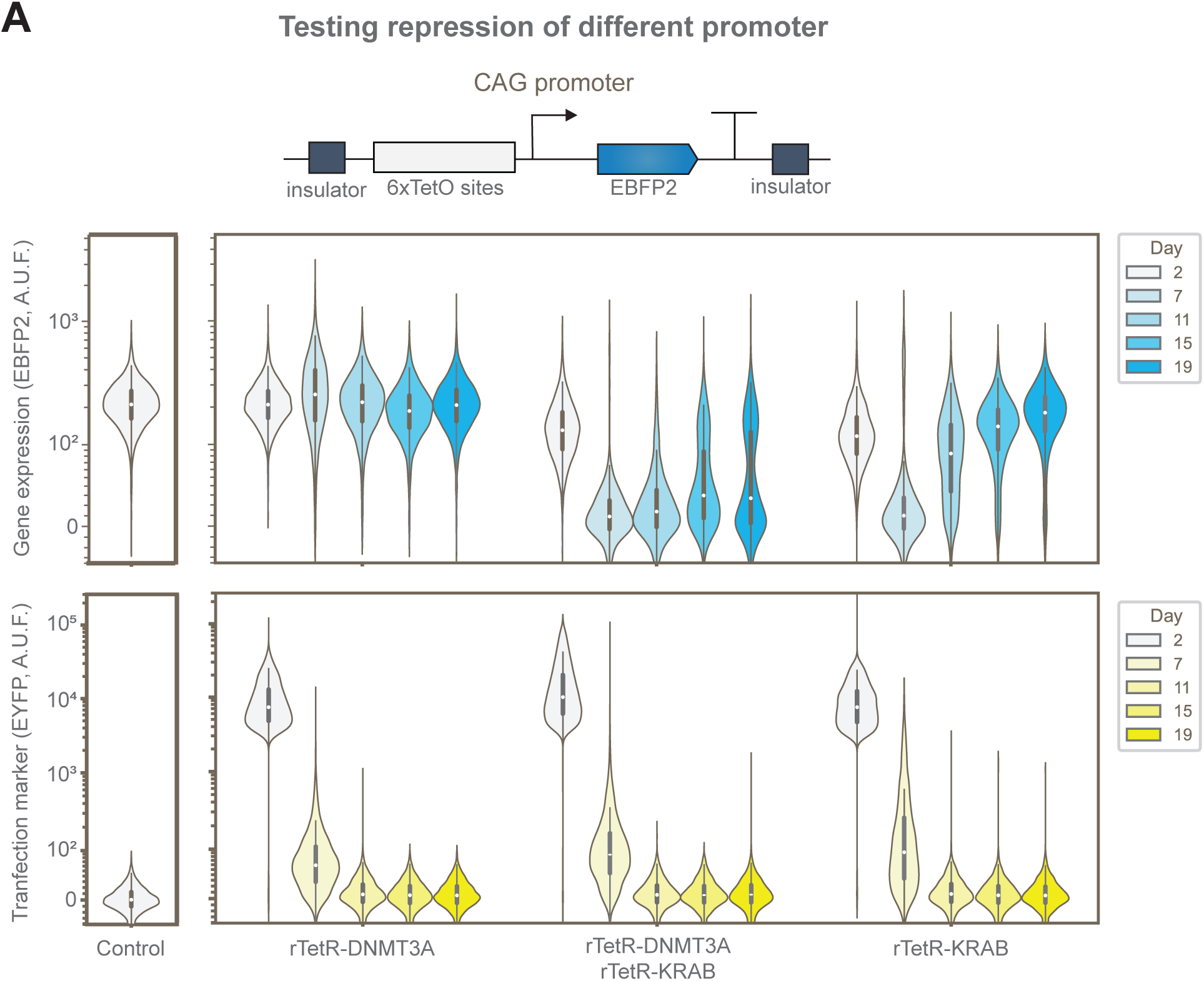
Induced silencing of the reported construct with CAG promoter in hiPSCs. **A.** Violin plots showing reporter gene expression (EBFP2) and transfection marker expression (EYFP) during silencing of the CAG promoter by transient transfection of rTetR-DNMT3a, rTetR-KRAB or both in combination. Data are plotted from two independent replicates. The white dot represents the median, the thick box represents the interquartile range (IQR) and the thin gray line represents 1.5x the IQR.

**Fig. S7.**
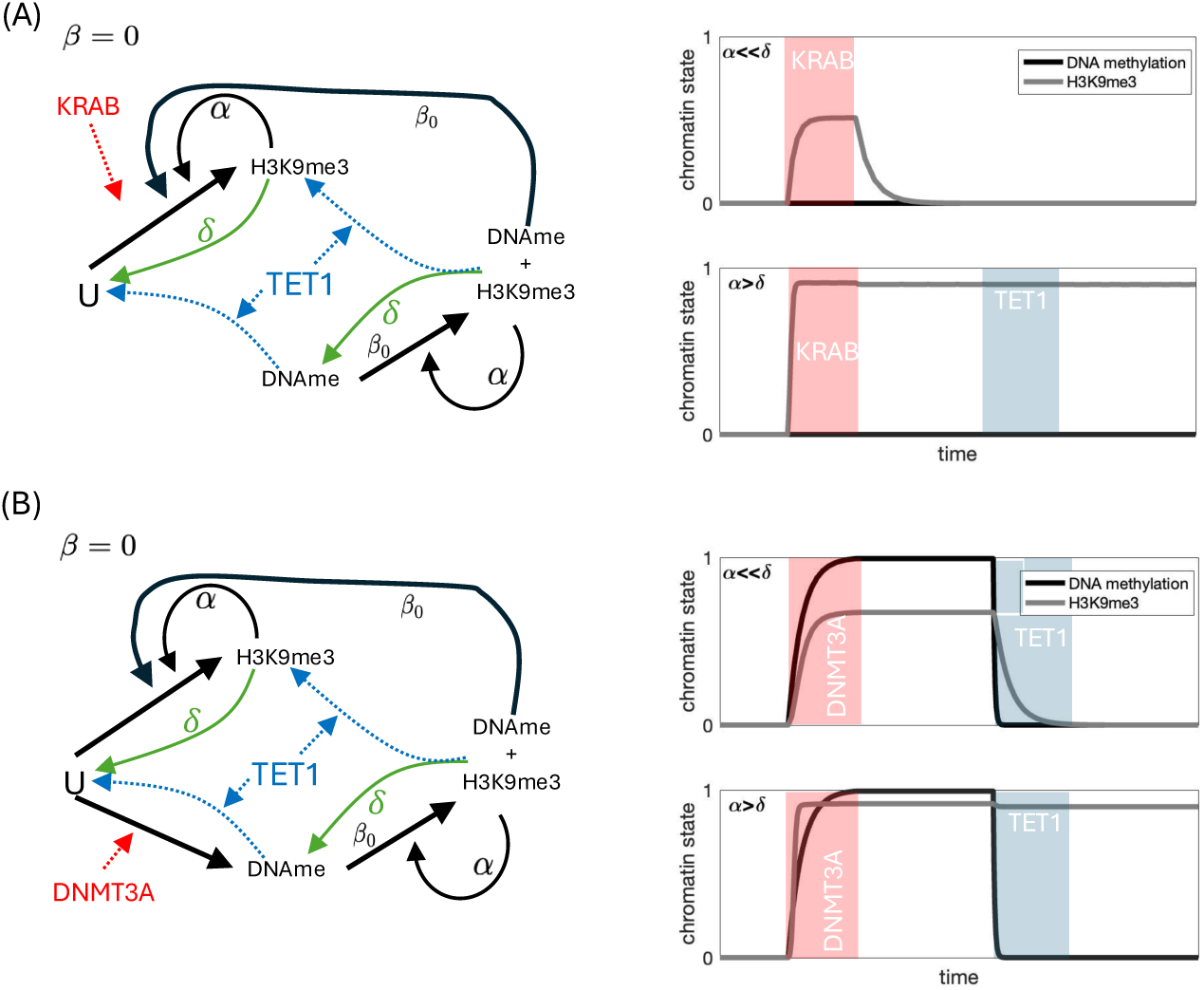
Chromatin modification circuit where H3K9me3 does not recruit writers of DNA methylation, that is, *β* = 0 **(CHO-K1 cells). A.** Transient recruitment of KRAB leads to establishing H3K9me3 only transiently when autocatalysis of H3K9me3 overpowers its decay (*α ≪ δ*) while it leads to stable H3K9me3 otherwise (*α > δ*). **B.** Recruitment of DNMT3A leads to stable H3K9me3 and DNA methylation independent of whether H3K9me3 autocatalysis overpowers its decay. Parameters: *β*_0_ = 1, *α* = *{*0.1, 10*}*, *δ* = 1, *T* = 10, *D* = 1, and *K* = 1.

**Fig. S8.**
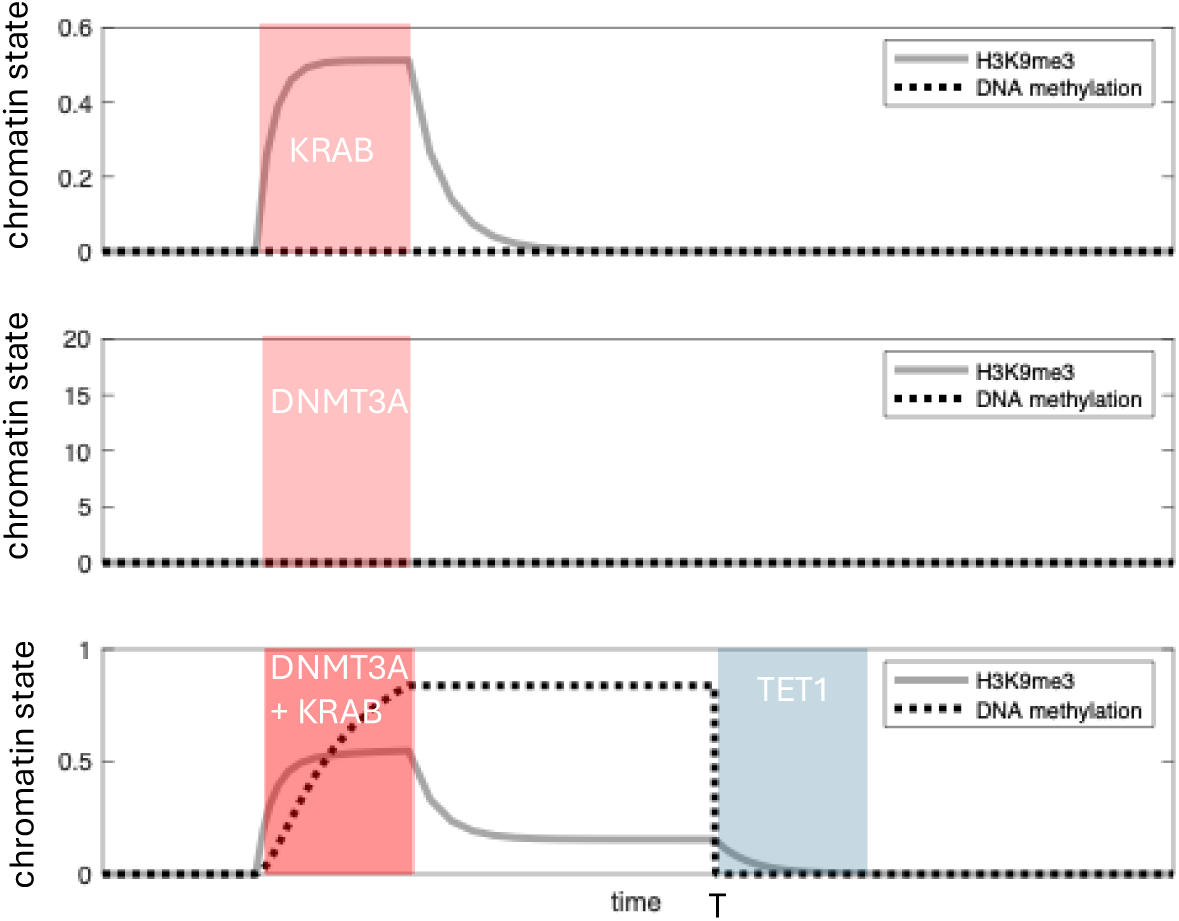
Simulation results when writing DNA methylation requires H3K9me3 to be present. Simulation results from the ODE model in SI Section 1, in which *α* = 0.1, *δ* = 1, *β* = *β*_0_ = 0.1, *T* = 10, *K* = *{*0, 1*}*, *D* = *{*0, 4*}*, and we have replaced *D · u* by zero, modeling that DNMT3A cannot write DNA methylation on nucleosomes devoid of H3K9me3, and we have replaced *β* by *β · D*, modeling that H3K9me3 can recruit DNA methylation only when we overexpress rTetR-DNMT3A.

**Fig. S9.**
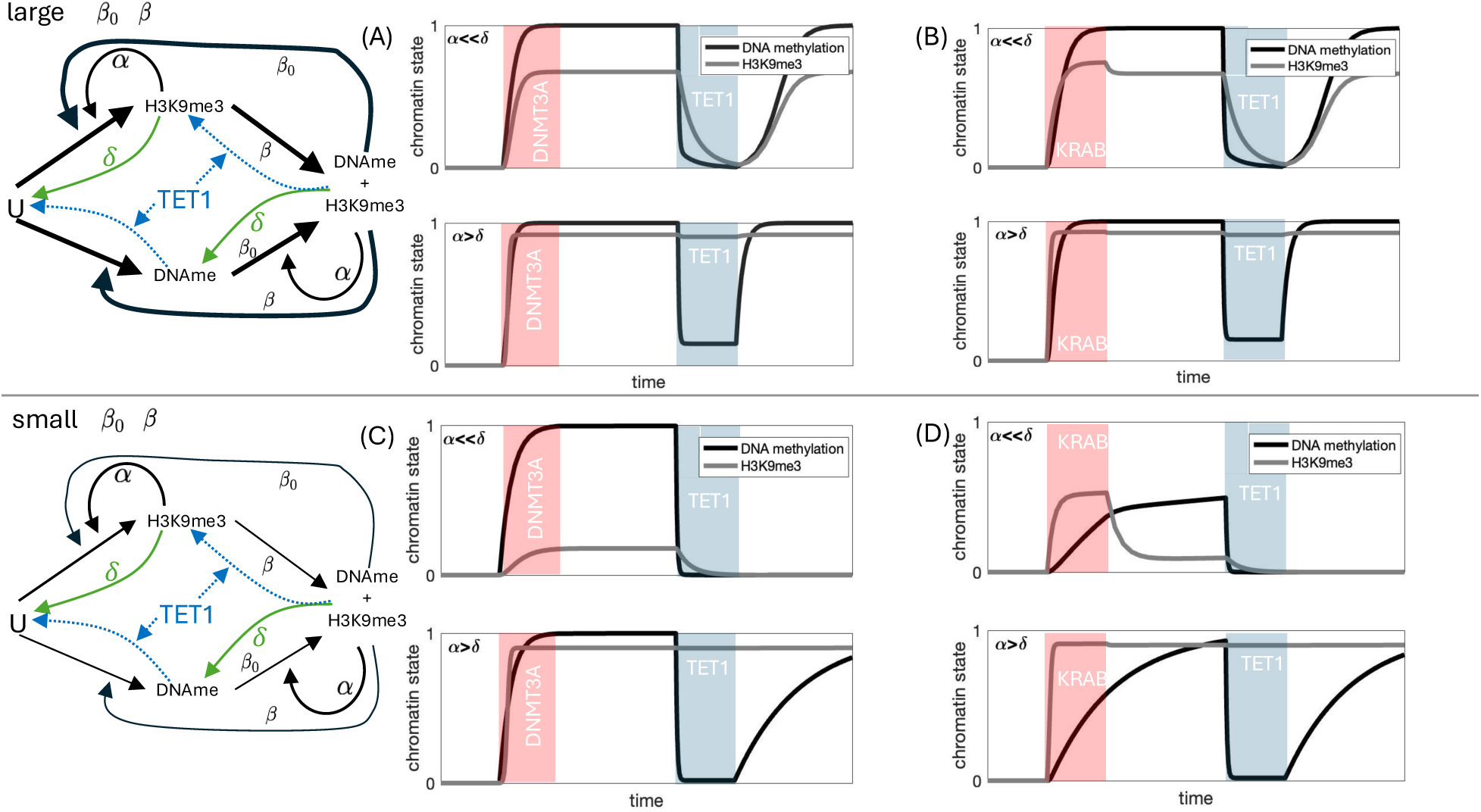
Effect of the strength of the positive feedback between DNA methylation and H3K9me3 on the ability of TET1 to permanently remove H3K9me3. A-B. When the positive feedback between DNA methylation and H3K9me3 is sufficiently strong (*β* and *β*_0_ are large), independent of whether the autocatalysis of H3K9me3 is strong (*α > δ*), after TET1 is removed, both marks reinforce each other and build back up. **C-D.** When the positive feedback between DNA methylation and H3K9me3 is sufficiently weak (*β* and *β*_0_ are large), DNA methylation and H3K9me3 will not build back up unless the autocatalysis of H3K9me3 is sufficiently strong (*α > δ*). Top: *β* = *β*_0_ = 1. Bottom: *β* = *β*_0_ = 0.1. Other parameter values: *α* = *{*0.1, 10*}*, *δ* = 1, *T* = 10, *D* = 1, and *K* = 1.

**Fig. S10.**
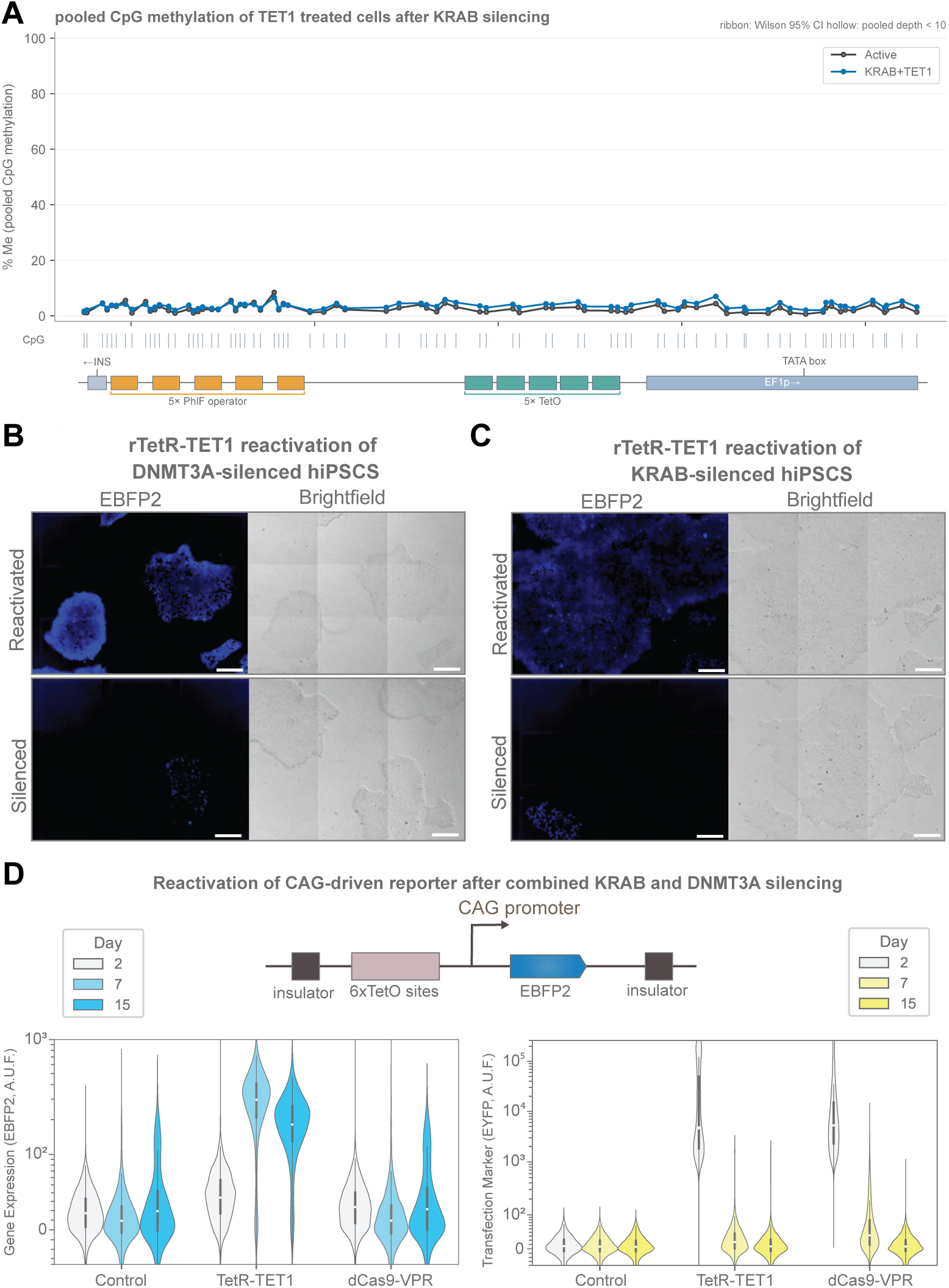
CpG line plot, single channel microscopy images and CAG promoter reactivation. **A.** Line plot showing CpG methylation levels of the transgene operator-promoter pair for cells in the active state and after TET1-mediated reactivation of KRAB-silenced cells. Data are average from three independent replicates and shaded error represents 95% confidence interval. **B-C.** Single channels for microscopy images of rTetR-TET1 reactivation experiment following **(B)** rTetR-DNMT3A silencing or **(C)** PhlF-KRAB silencing. Scale bars are 300 *µ*m. **D.** Violin plots showing TetR-TET1 mediated reactivation dynamics of CAG-driven transgene reporter in hiPSCs following silencing PhlF-KRAB and rTetR-DNMT3A co-transfections. Data are plotted from three independent replicates. The white dot represents the median, the thick box represents the interquartile range (IQR) and the thin gray line represents 1.5x the IQR.

**Fig. S11.**
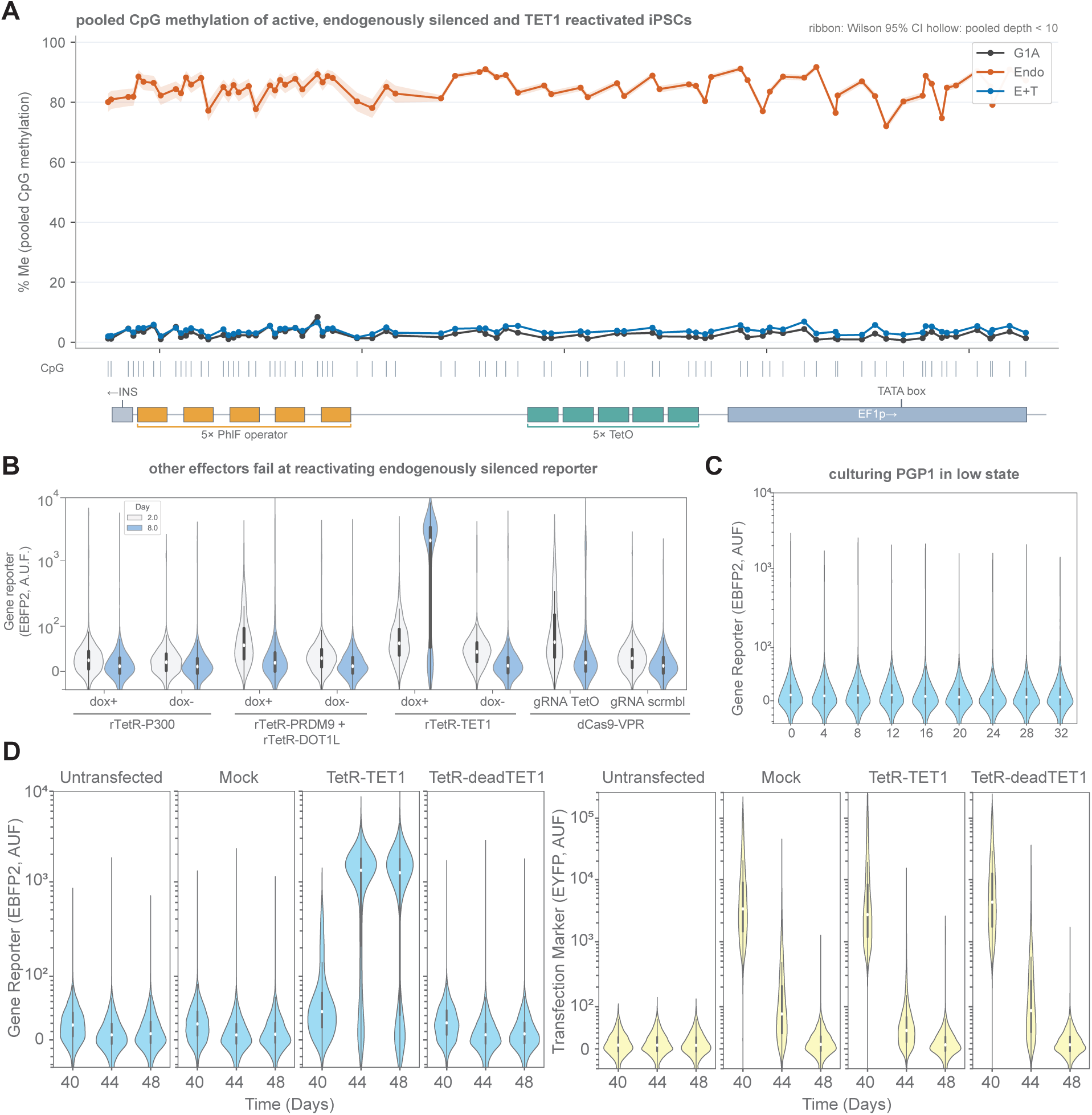
Single channel microscopy images and RNA seq data. **A.** Line plot showing CpG methylation levels of the transgene operator-promoter pair for cells in the active state, cells that have been endogenously silenced, and after TET1-mediated reactivation. Data are average from three independent replicates and shaded error represents 95% confidence interval. **B.** Violin plots showing reporter gene expression (EBFP2) of hiPSCs 4 days after transfection with different chromatin regulators and the VP64-P65-Rta (VPR) transcriptional activator. **C.** Time-resolved violin plots showing reporter gene expression (EBFP2) of endogenously silenced hiPSCs cultured int he low state for 32 days. **D.** Violin plots showing reporter gene expression (EBFP2) and transfection marker expression (EYFP) of the long term culture of endogenously silenced hiPS cells after reactivation by transient TET1 transfection along with mock controls (plasmid backbone) and catalytically inactive TET1 (deadTET1). For all violin plots, data are plotted from three independent replicates and the white dot represents the median, the thick box represents the interquartile range (IQR) and the thin gray line represents 1.5x the IQR.

**Fig. S12.**
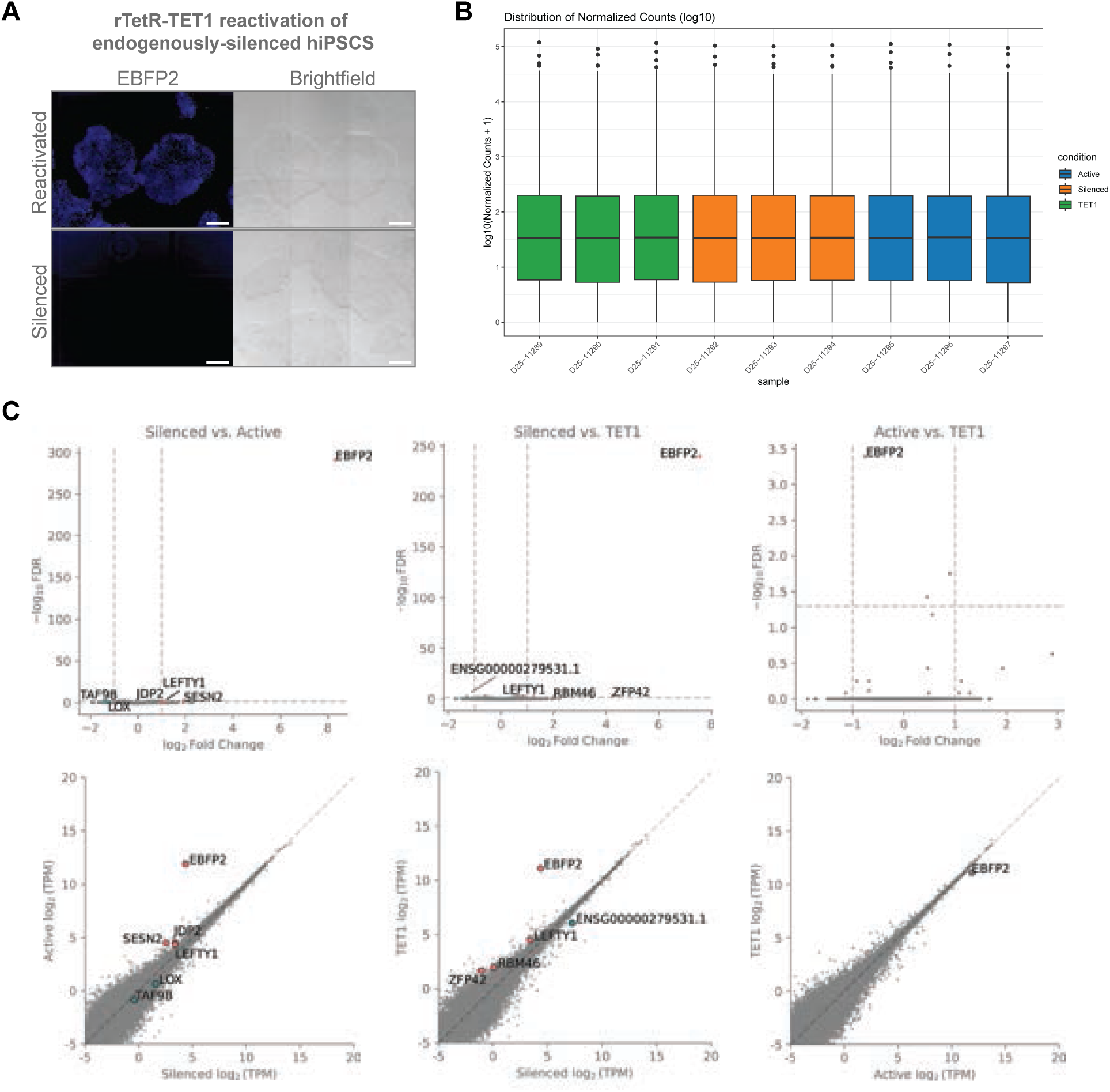
Single channel microscopy images and RNA seq data. **A.** Single channels for microscopy images of rTetR-TET1 reactivation experiment following endogenous silencing. This panel is related to Fig. 5E. Scale bars are 300 *µ*m. **B.** Normalized transcript counts for each replicate in RNA seq experiment. This panel are related to Fig. 5G and the following panels. **C.** Volcano plots and transcript counts of RNA seq data for the different conditions. These panels are related to Fig. 5G. TPM: Transcripts per million; FDR: False discovery rate.

**Fig. S13.**
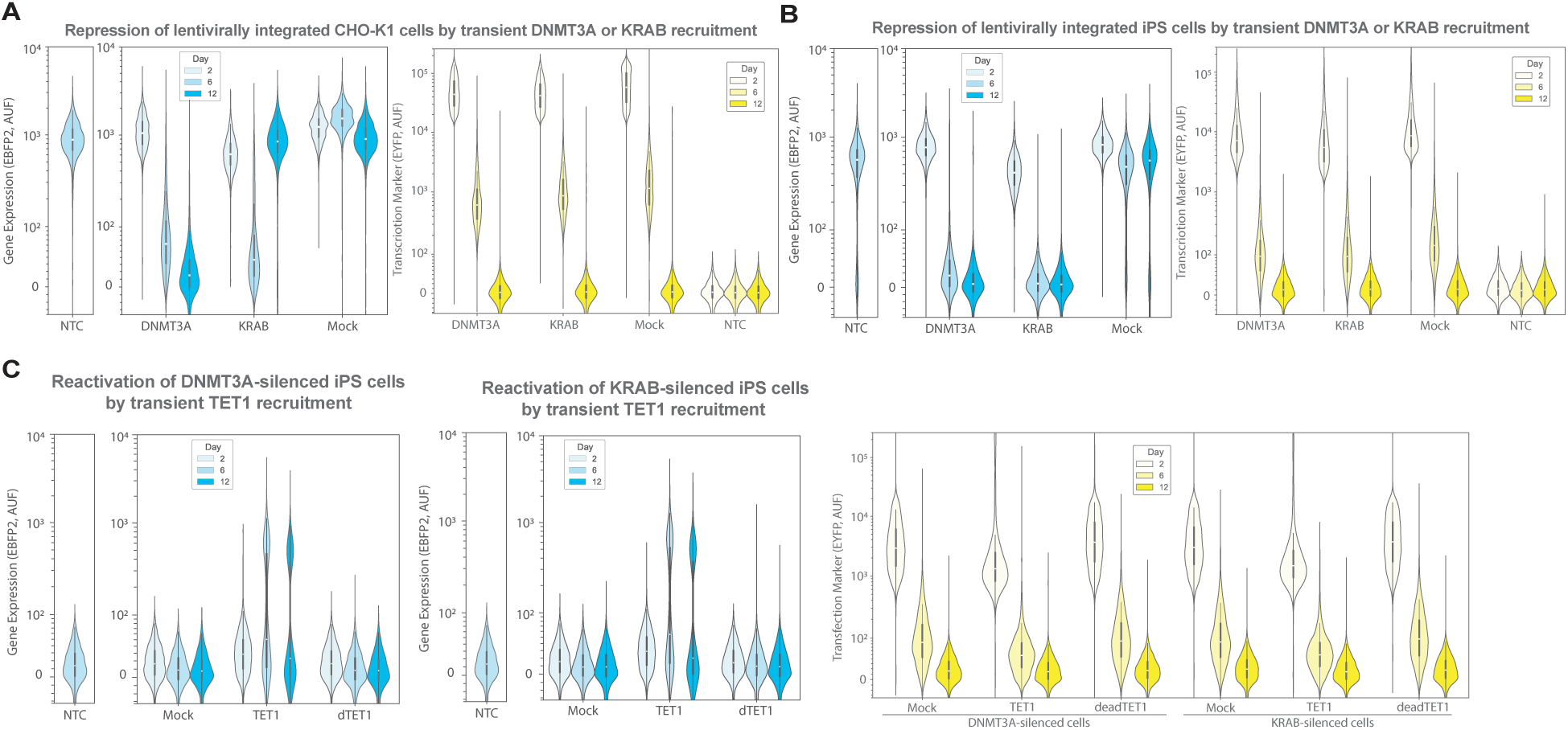
Lentiviral reporter gene regulation by targeted chromatin editing. **A.** Violin plots showing lentiviral reporter gene expression (EBFP2) and transfection marker expression of CHO-K1 cells after repression by targeted DNMT3A or KRAB treatment, along with mock control (plasmid backbone). **B.** Violin plots showing lentiviral reporter gene expression (EBFP2) and transfection marker expression of hiPS cells after repression by targeted DNMT3A or KRAB treatment, along with mock control (plasmid backbone). **C.** Violin plots showing lentiviral reporter gene expression (EBFP2) and transfection marker expression (EYFP) of DNMT3A- or KRAB-silenced hiPS cells after reactivation by transient TET1 transfection along with mock controls (plasmid backbone) and catalytically inactive TET1 (dTET1). For all violin plots, data are plotted from three independent replicates and the white dot represents the median, the thick box represents the interquartile range (IQR) and the thin gray line represents 1.5x the IQR.

